# Replication timing maintains the global epigenetic state in human cells

**DOI:** 10.1101/2019.12.28.890020

**Authors:** Kyle N. Klein, Peiyao A. Zhao, Xiaowen Lyu, Daniel A. Bartlett, Amar Singh, Ipek Tasan, Lotte P. Watts, Shin-ichiro Hiraga, Toyoaki Natsume, Xuemeng Zhou, Danny Leung, Masato T. Kanemaki, Anne D. Donaldson, Huimin Zhao, Stephen Dalton, Victor G. Corces, David M. Gilbert

## Abstract

DNA is replicated in a defined temporal order termed the replication timing (RT) program. RT is spatially segregated in the nucleus with early/late replication corresponding to Hi-C A/B chromatin compartments, respectively. Early replication is also associated with active histone modifications and transcriptional permissiveness. However, the mechanistic interplay between RT, chromatin state, and genome compartmentalization is largely unknown. Here we report that RT is central to epigenome maintenance and compartmentalization in both human embryonic stem cells (hESCs) and cancer cell line HCT116. Knockout (KO) of the conserved RT control factor RIF1, rather than causing discrete RT switches as previously suspected, lead to dramatically increased cell to cell heterogeneity of RT genome wide, despite RIF1’s enrichment in late replicating chromatin. RIF1 KO hESCs have a nearly random RT program, unlike all prior RIF1 KO cells, including HCT116, which show localized alterations. Regions that retain RT, which are prevalent in HCT116 but rare in hESCs, consist of large H3K9me3 domains revealing two independent mechanisms of RT regulation that are used to different extents in different cell types. RIF1 KO results in a striking genome wide downregulation of H3K27ac peaks and enrichment of H3K9me3 at large domains that remain late replicating, while H3K27me3 and H3K4me3 are re-distributed genome wide in a cell type specific manner. These histone modification changes coincided with global reorganization of genome compartments, transcription changes and a genome wide strengthening of TAD structures. Inducible degradation of RIF1 revealed that disruption of RT is upstream of genome compartmentalization changes. Our findings demonstrate that disruption of RT leads to widespread epigenetic mis-regulation, supporting previously speculative models in which the timing of chromatin assembly at the replication fork plays a key role in maintaining the global epigenetic state, which in turn drives genome architecture.

## Main Text

DNA is replicated during S phase of the cell cycle in a temporal order known as the replication timing (RT) program. RT is conserved among eukaryotes, cell type specific and correlates with many important epigenomic features (*1*). Early replicating chromatin has euchromatic characteristics such as active histone modifications and location in the nuclear interior while late replicating chromatin is associated with heterochromatic features like transcriptionally repressive histone modifications and location at the nuclear periphery. Early and late replicating chromatin correspond to A- and B-compartments respectively as defined by high throughput chromatin conformation capture (Hi-C) (*2*). Despite these close correlations, the mechanistic link between RT and the accurate maintenance of chromatin through cell cycles remains elusive. Prior work has shown that histones and their modifications are both recycled from parental chromatin and added and modified de novo after passage of the replication fork with different chromatin states showing differing dynamics of reassembly (*3, 4*). It has long been hypothesized that RT influences chromatin maintenance. Indeed, microinjection of plasmids into mammalian nuclei revealed that plasmids replicated in early S phase were decorated with acetylated histones, while those replicated later in S phase were devoid of acetylated histones (*5*). However, there is still no direct evidence implicating RT in epigenetic state maintenance, largely due to the inability to manipulate genome wide RT. Recently the conserved protein RIF1 has been shown to affect RT in many eukaryotes, however, partly because the effects have been partial or localized, RIF1 disruption has not been exploited to study the effects of RT abrogation (*6*–*11*).

To gain insight into the role of genome wide RT as controlled by RIF1 in shaping the epigenome we examined the effects of RIF1 knock out (KO) in three human cell lines. H9 hESC, HCT116, established by removal of RIF1 exon 3 (Fig. S1a, b, c). As previously reported (*11*), RIF1 KO cells proceeded through the cell cycle with minor decreases in the number of S phase cells and increases in the number of gap phase cells (Fig S1d, e). All three RIF1 KO cell lines exhibit genome wide aberrations in RT measured using E/L Repli-seq (*12*) but with varying degrees of severity (Fig. 1a). Similar to all prior reports in mammalian cells (*8, 9, 11*), discrete domains changed RT either from early to late (EtL) or late to early (LtE) in E/L RT profiles of HCT116 and HAP1 cells. RIF1 KO caused 43% of the genome to change RT (23% EtL and 20% LtE) in HCT116 and HAP1 cells. However, in H9 hESCs (Fig. 1a), nearly the entire genome acquired a log2E/L dynamic range close to zero (Fig 1b), precluding identification of specific domain level RT changes and suggesting that RIF1 is necessary for nearly all temporal control of replication in hESCs. This near complete loss of RT control in RIF1 KO hESCs was much more severe than we previously reported in RIF1 KO mouse ESCs (*11*) (Fig S1f). Moreover, partial knockdown (KD) of RIF1 in H9 hESCs using an shRNA (Fig S2a) resulted in a partial effect on the RT program (Fig S2b), demonstrating that RIF1’s control of global RT in hESCs is dosage dependent. Plotting the density of RT values as a histogram confirmed that all three RIF1 KO cell lines, as well as RIF1 KD H9 hESCs, show genome wide RT values accumulating near the middle of S phase for most genomic bins (Fig 1b, Fig S2c) and that H9 RIF1 KO cells showed a sharp genome wide peak of log2E/L values at zero (Fig 1b). Replication foci patterns assayed by BrdU incorporation, which track with genome wide RT (*13*) (Fig S3a), were also altered (Fig S3b) as RIF1 KO cells showed a loss of clear middle S phase patterns and ‘blending’ of early and middle replication foci patterns but maintained distinct early and late foci patterns (Fig S3a, b). Together, these results demonstrate a considerably more extensive role for RIF1 in RT control in hESCs than in any other mammalian cell type so far studied.

**Figure 1.**
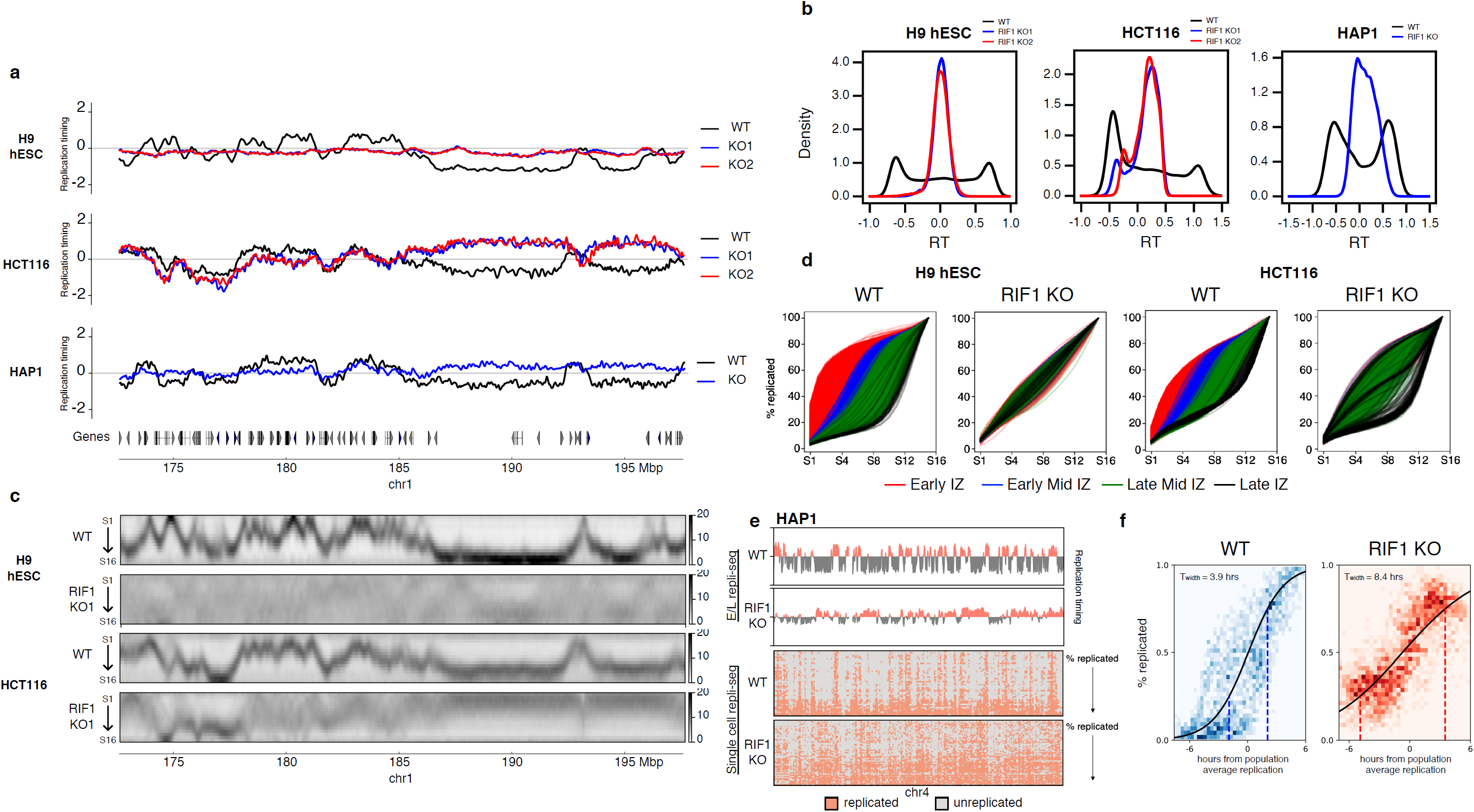
RIF1 controls RT by reducing cell to cell variation in replication timing. a, E/L Repli-seq plots of Chr1 172.6-197.6 Mb in WT and two RIF1 KO clones overlaid in H9 hESCs (top) and HCT116 (middle) and HAP1 (bottom) cell lines. b, Genome wide probability density of E/L RT values in H9 hESCs (left), HCT116 (middle), and HAP1 (right) WT (black) and RIF1 KO (blue and red). c, High resolution Repli-seq plots of Chr1 172.6-197.6 Mb in WT and RIF1 KO in H9 hESCs (top two) and HCT116 (bottom two); same locus as a. d, Cumulative percent-replicated plots for each IZ called in WT cells versus S phase fraction of 16 fraction Repli-seq color coded by their timing (red: early, blue: early mid, green: late mid, black: late). e, Binarized single-cell Repli-seq heatmaps of Chr4 in HAP1 WT (top) and RIF1 KO (bottom) sorted by percentage replicated where the top row represents the cell with the lowest percentage of genome replicated and the last row represents the cell with the most percentage of genome replicated. f, Sigmoidal fitting of the percentage replicated against time in hours from population average replication for HAP1 WT (left) and RIF1 KO (right). The heatmaps (blue for WT and red for KO) represent the data spread of percentage replicated (y-axis) against time from population average replication (x-axis) for all 50kb bins genome wide for all cells. Dotted lines at 25% of cells replicated and 75% of cells replicated indicate the span of T_width_.

Previous reports have interpreted RT changes in RIF1 KO cells as distinct RT switches (*8, 9, 11*) but the extent of the RT phenotype in RIF1 KO hESCs made us hypothesize that RIF1 KO results in a loss of temporal replication specificity resulting from increased cell to cell RT heterogeneity within the population. To address this hypothesis, we applied our recently developed high resolution Repli-seq protocol (*14*), which uses a shorter nascent DNA labelling period and divides S phase into 16 small, evenly distributed fractions. High resolution Repli-seq produces RT heatmaps that capture peaks of replication initiation termed initiation zones (IZs) and large valleys of late replication that are sites of termination containing broadly distributed, low efficiency initiation events (*14*). Distinct changes in RT, such as those that occur during stem cell differentiation, manifest as clear EtL or LtE shifts of peaks and valleys on the temporal axis (*14*), whereas greater RT variation would manifest as highly diffused peaks or valleys resulting from reads spread out across many S phase fractions for any genomic location. High resolution Repli-seq of RIF1 KO cells revealed dramatic diffusion of RT patterns and loss of defined IZs in both HCT116 and H9 hESCs (Fig 1c, Fig S4a, b) indicating major RT variation within the cell population. EtL and LtE regions in RIF1 KO HCT116 cells called in E/L Repli-seq showed major loss of temporal control and replication across a broad distribution of times during S phase in the high resolution Repli-seq (Fig S4c). Surprisingly, even early replicating regions that were not called as statistically confident EtL switches in the HCT116 E/L Repli-seq also lost definition of IZs and RT control when assayed using the more sensitive high resolution Repli-seq (Compare Figs. 1a, c, Fig S4c). Thus, the entirety of the early replicating genome lost RT control upon RIF1 KO in both cell lines. By contrast, many late replicating regions in HCT116 not called as LtE switches in E/L Repli-seq retained late replication control when assayed by high resolution Repli-seq (Fig S4c) indicating a separate, RIF1-independent mechanism controlling RT for these late regions in HCT116, which we will expand upon below. Remarkably, high resolution Repli-seq of RIF1 KO H9 hESCs resulted in a heatmap that lacked distinct IZs and valleys, suggesting a near complete genome-wide loss of temporal replication control in which any sequence can replication at any time in the cell population (Fig 1d).

To better quantify the extent and temporal direction of the loss of RT control, we calculated RT indices for all genomic bins of high resolution Repli-seq in WT and RIF1 KO samples where positive values indicate early replication and negative values indicate late replication (Fig S4d, Methods). Again, early replicating regions that were not called as EtL switches in the HCT116 E/L Repli-seq showed noticeable changes in their RT indices similar to called EtL switches (Fig S4e). RT differences (RT_diff_) were then calculated by subtracting RIF1 KO RT indices from WT RT indices (Methods). We further applied a 3 component Gaussian Mixture model to RT_diff_ distribution to identify genomic bins that were characterized by: 1) negative RT_diff_, which are regions that are later replicating in WT and earlier replicating in KO, 2) positive RT_diff_, which are regions that are earlier replicating in WT and later replicating in KO, 3) close-to-zero RT_diff_, which are regions that showed limited RT difference between WT and KO or could not be called as significantly different (Fig S4f, Methods). 86% and 78% of the genome showed significant RT_diff_ in H9 and HCT116, respectively (Fig S4f table). These results demonstrate a much more significant effect of RIF1 loss on RT in both cell types than previously imagined.

To examine the effect of RIF1 deletion on replication initiation within IZs genome wide, we divided each IZ called in high resolution WT Repli-seq data into timing categories based on the temporal position of the IZ peak in S phase (*14*) and plotted the cumulative percentage of DNA replicated through S phase in both WT and RIF1 KO cells (Fig 1d). WT cells showed typical segregation of IZs according to the temporal order associated with steady state replication and steep sigmoidal-like curves (*14*) while RIF1 KO cells showed major overlap of IZ classes and flatter sigmoidal-like curves (Fig 1d) showing that RIF1 KO caused loss of genome wide replication initiation timing specificity in both cell lines, with H9 hESCs losing any detectable temporal control. To quantify this change we calculated the parameter T_width_ as the time between when 25% and 75% of cells replicate for each genomic bin. A small T_width_ is indicative of a more deterministic replication scheme whereas a larger T_width_ is representative of greater heterogeneity. Genome wide measurement of T_width_ greatly increased in both cell types upon RIF1 KO indicating a major increase in RT heterogeneity (Fig S4g).

To directly validate that the RT aberration seen in RIF1 KO was due to increased cell to cell RT heterogeneity we performed our recently developed single-cell Repli-seq (*15*) on human haploid HAP1 WT and RIF1 KO cells, which eliminate allelic RT variation to enable single cell analysis. RIF1 KO caused a major increase in RT heterogeneity genome wide, confirming this effect in a third cell line. We ranked binarized single cell RT profiles by their percentage of genome replicated indicating their progression through S phase and constructed heatmaps representing replicated or unreplicated windows for WT and RIF1 KO cells (Fig 1e). WT heatmaps correspond well to E/L Repli-seq, early replicating regions manifested as distinct domains that have finished replication early in S phase in the majority of cells whereas late replicating regions remained unreplicated until the later stages of S phase (Fig 1e). In KO heatmaps, however, the distinction between early and late replicating domains was blurred at all stages of S phase (Fig 1e, Fig S4h). To quantify this heterogeneity genome wide we assumed a 10 hour S phase and calculated ‘time from population average replication according to E/L RT values’ at 0.1 intervals for each single cell (*15*) and plotted the relationship between percentage of genomic bins replicated to their ‘time from population average replication’ in the form of heatmaps (Fig 1f). We then performed sigmoidal fitting on the heatmap and calculated the parameter T_width_ for each bin (See Methods). HAP1 WT cells’ T_width_ was 3.9 hours while the absence of RIF1 greatly increased the T_width_ to 8.4 hours indicating a major increase in RT heterogeneity (Fig 1f). Taken together, both high resolution Repli-seq and single cell Repli-seq demonstrate that loss of RIF1 in 3 human cell lines disrupts RT by dramatically increasing temporal heterogeneity of replication across the cell population rather than causing discrete RT shifts in all cells.

Mouse RIF1 was previously reported to bind late replicating chromatin (*11*). We performed Cut&Run (*16*) against GFP on GFP tagged RIF1 in HCT116 and H9 hESCs (Fig S1a, b) to map RIF1 binding in human cells. Consistent with mESCs (*11*), RIF1 binding was enriched in the late replicating portion of the genome in both cell lines (Fig S5a) and bound chromatin in broad domains rather than distinct peaks (Fig 2a). We first divided the late replicating genome into regions that lost RT control upon RIF1 KO, characterized by continuous bins displaying negative RT_diff_ (Fig 2b blue in WT, red in KO) and those that maintained their late RT upon RIF1 KO, characterized by continuous bins displaying close-to-zero RT_diff_ (Fig 2d blue in WT, blue in KO). These regions are hereafter referred to as ‘affected’ or ‘unaffected’ late regions respectively (Fig 2a, b, c, d). RIF1 binding was especially enriched at affected domains (Fig 2a, e) while unaffected domains showed lower RIF1 enrichment in both cell lines (Fig 2c, e). Contrary to what was reported in mESCs (*11*), unaffected regions showed no preferential association with the nuclear lamina than affected regions via Lamin B1 DamID (Fig 2b, c).

**Figure 2.**
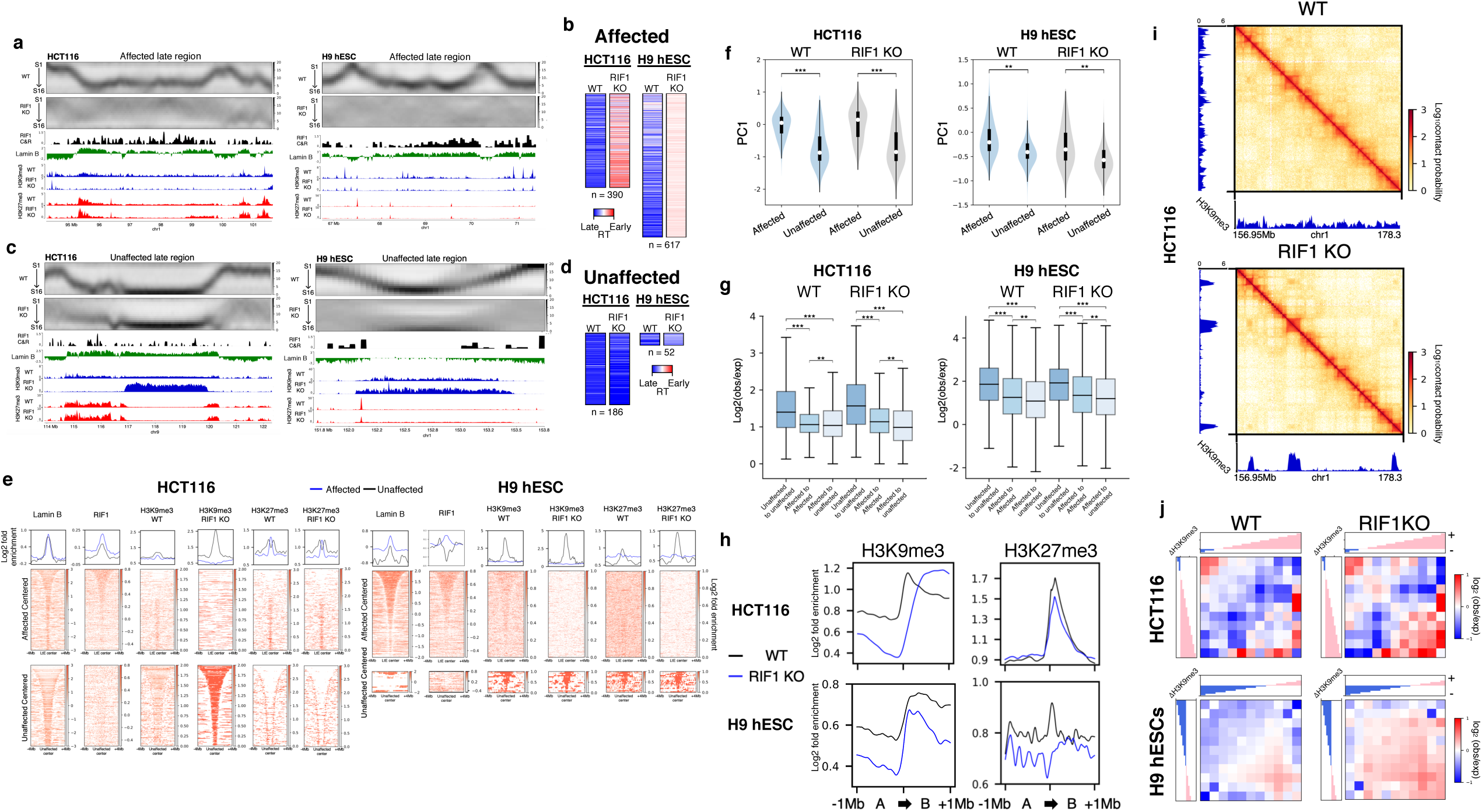
RT affected and unaffected regions are distinct classes of chromatin. a, Affected region at Chr1 94.3-101.6 Mb in HCT116 (left) and Chr1 66.75-71.4 Mb in H9 hESCs (right) showing from top to bottom: 16 fraction Repli-seq in WT and RIF1 KO cells, RIF1 Cut&Run in WT cells, Lamin B1 DamID in WT cells, H3K9me3 ChIP-seq in WT and RIF1 KO cells, and H3K27me3 ChIP-seq in WT and RIF1 KO cells. b, Heat maps of RT indices for affected regions in WT and RIF1 KO of HCT116 and H9 hESCs. c, Unaffected region at Chr9 113.95-122.3 Mb in HCT116 (left) and Chr1 151.7-153.75 Mb in H9 hESCs (right) showing the same panels as a. d, Heat maps of RT indices for unaffected regions in WT and RIF1 KO of HCT116 and H9 hESCs. e, Fold enrichment signal pile ups of signal from indicated assay in HCT116 (left) and H9 hESC (right) WT or RIF1 KO cells centered on affected regions (top) or unaffected regions (bottom) ±4 Mb and sorted by size. Line plots represent cumulative signal in WT (black) and RIF1 KO cells (blue). f, Violin plots of PC1 Eigenvector values for LtE and unaffected chromatin loci in WT and RIF1 KO HCT116 (left) and H9 hESCs (right); (***p < 0.0005, **p < 0.005). g, Box plots of interaction strength between and within LtE and unaffected chromatin domains in WT and RIF1 KO HCT116 (left) and H9 hESCs (right); (***p < 0.0005, **p < 0.005). h, Pile up line plots of indicated histone marks at A to B compartment boundary ±1 Mb in WT (black) and RIF1 KO (blue) HCT116 and H9 hESCs. i, H9K9e3 ChIP-seq tracks beside ICE normalized Hi-C contact maps of HCT116 WT and RIF1 KO. j, Log2(obs/exp) aggregate interactions between B compartments in WT and RIF1 KO HCT116 (top) and H9 hESCs (bottom), The interactions were binned into 11 equal segments, which were ranked by increasing delta H3K9me3 within B compartments where negative and positive values indicate decrease and increase in H3K9me3 between WT and RIF1KO.

To further investigate these late replicating regions, we performed ChIP-seq on the late replication associated marks H3K9me3 and H3K27me3 in both HCT116 and H9 hESC RIF1 KO cells. H3K9me3 marked regions manifest as either small peaks or large megabase scale domains (*17*). In WT cells, affected regions were enriched for smaller H3K9me3 peaks while unaffected regions contained larger H3K9me3 domains (Fig 2a, c, e, Fig S5b). RIF1 KO caused genome wide changes to H3K9me3 peaks and domains in both cell lines (Fig S5c). Without RIF1, small H3K9me3 peaks were lost, coincident with a loss in RT control (Fig 2a, c, e). However, the large H3K9me3 domains, which are far more abundant in HCT116 than H9 hESCs, dramatically increased in their density of H3K9me3 and retained their late RT in the absence of RIF1 (Fig 2c, e). In fact, almost all regions that retained late RT in RIF1 KO HCT116 cells (n=209) were large H3K9me3 domains. The rare domains that retained late RT in H9 hESCs (n=49) were also large H3K9me3 domains. Overall levels of H3K9me3 within RIF1 KO H9 hESCs did not change compared to WT (Fig S5d) suggesting that total H3K9me3 is redistributed rather than gained or lost upon RIF1 KO. Most H3K9me3 unaffected regions do not overlap between cell types, consistent with the model that these H3K9me3 domains are developmentally regulated and implicated in silencing of lineage inappropriate genes (*17*) (Fig S5e). By contrast, H3K27me3 changes after RIF1 KO were cell type specific (Fig 2a, c, e). H3K27me3 peaks were enriched at affected regions but depleted from unaffected regions in WT HCT116 cells and these distributions were unaffected by RIF1 KO (Fig 2a, c, e). However, H3K27me3 was similar to H3K9me3 in H9 hESCs in that RIF1 loss resulted in the depletion of H3K27me3 peaks at affected regions and enrichment at the few unaffected regions (Fig 2a, c, e). Thus, RIF1 and H3K9me3 domains are two separate mechanisms that orchestrate late RT to different extents in different cell types.

The strong association between late replication and B compartmentalization as defined by negative PC1 Eigenvector values (*18, 19*) compelled us to investigate the genomic compartmentalization and architecture of RIF1 KO cells by Hi-C (*20*). Surprisingly, we found that affected and unaffected regions, despite being similarly late replicating, have distinct compartmental identities. In WT cells unaffected regions were associated with more negative PC1 Eigenvector values and thus interacted more strongly with other B compartment regions than affected regions (Fig 2f), while affected regions showed a slight interaction preference to the A compartment particularly in HCT116 cells (Fig 2f). Upon RIF1 KO, the interaction between unaffected regions with the rest of the B compartment was strengthened whereas affected regions display more preferential interactions with the A compartment (Fig 2f) indicating RIF1 delays the replication of regions in A compartment that would otherwise be earlier replicating. Moreover, WT interactions between unaffected regions were significantly stronger than interactions between affected regions or interactions between unaffected regions and affected regions (Fig 2g) further confirming that affected and unaffected regions are two intrinsically different classes of late replicating regions with distinct interaction preferences. These distinct interaction preferences further increase in RIF1 KO (Fig 2g). We next looked to see if RIF1 KO had an effect on the distribution of histone marks between genomic compartments by plotting the aggregate log2 fold enrichment of histone modification ChIP-seq centered at the A/B compartment boundary (Fig 2h). H3K9me3 was depleted from the A compartment and enriched in the B compartment in HCT116 RIF1 KO cells (Fig 2h) consistent with the fact that in HCT116 affected regions were more associated with the A compartment while unaffected regions were predominately within the B compartment (Fig 2f). Similarly, RIF1 KO H9 hESCs had depleted H3K9me3 in the A compartment, however the B compartment as a whole also lost H3K9me3 (Fig 2h) as the majority of B compartment regions were affected and lost late replication specificity. H3K27me3 changes were cell type specific as distributions remained largely unchanged in HCT116 but showed a decrease in both compartments in H9 hESCs (Fig 2h). These data indicate RT affected and unaffected regions are two distinct classes of late replicating chromatin which are differentially affected by RIF1 KO.

New interactions evident in Hi-C contact maps were formed between unaffected regions that became highly enriched for H3K9me3 in RIF1 KO cells (Fig 2i). Interactions between H3K9me3 peaks were strengthened after RIF1 KO and driven by enriched H3K9me3 domains in unaffected regions (Fig S5f, g). We further sorted and binned intra B-compartment interactions according to the extent of H3K9me3 peak changes (negative: downregulated, positive: upregulated). In WT cells upregulated H3K9me3 domains and downregulated H3K9me3 peaks form separate interaction hubs within the B compartment (Fig 2j left). Interactions within these hubs, particularly the upregulated H3K9me3 domains, strengthen dramatically upon RIF1 KO (Fig 2j right). These results demonstrate that H3K9me3 domains form strong interactions with one another constituting one type of B compartment that maintains its late RT and strengthens its interactions without RIF1 whereas regions that lack large H3K9me3 domains form a separate hub of interactions and require RIF1 to enforce late replication.

RIF1 KO and subsequent loss of global RT not only caused substantial changes in B compartment associated chromatin modifications and interactions, but it also had a genome-wide effect on A compartment associated chromatin modifications. ChIP-seq on H3K27ac and H3K4me3 revealed a genome wide reduction of H3K27ac peaks (Fig S5h) along with specific changes in H3K4me3 in RIF1 KO cells. Similar to H3K9me3, RIF1 KO did not cause a change in global levels of H3K27ac in H9 hESCs compared to WT (Fig S5i) again suggesting redistribution rather than global reduction of chromatin marks upon RIF1 KO. H3K27ac and H3K4me3 became majorly depleted in the A compartment of both cell lines with mild increases in their association with the B compartment (Fig 3a). Loss of active marks from the A compartment, particularly H3K27ac, was concurrent with changes in compartmentalization and Hi-C contacts in both cell lines. In H9 hESCs the majority of compartment changes were the disappearance of small A compartments into neighboring B compartments (Fig 3b, c), while HCT116 exhibited discrete shifts in compartmentalization in both directions from A to B as well as B to A (Fig 3b). In both cell lines the compartmental switches lead to a more consolidated appearance in the correlation matrix heatmap (Fig 3b). Loss of A compartment interactions in Hi-C contact maps corresponded to reduced H3K27ac peaks in both cell lines (Fig 3c (arrows)). Within all A to B compartment switches genome wide the level of H3K27ac was majorly depleted in both cell lines (Fig 3d). Genome-wide interaction pileups showed specific interactions between H3K27ac peaks and all other H3K27ac peaks were drastically reduced at A to B compartment switches (Fig 3e). Changes in genome organization were not due to changes in cell cycle distributions as RIF1 KD H9 hESCs exhibited similar changes to cell cycle as KO (Fig S1d, e) without changes in genome organization (Fig S2d, e, f). These data strongly suggest that depletion of active histone modifications, particularly H3K27ac, leads to loss of A compartment interactions in RIF1 KO cells.

**Figure 3.**
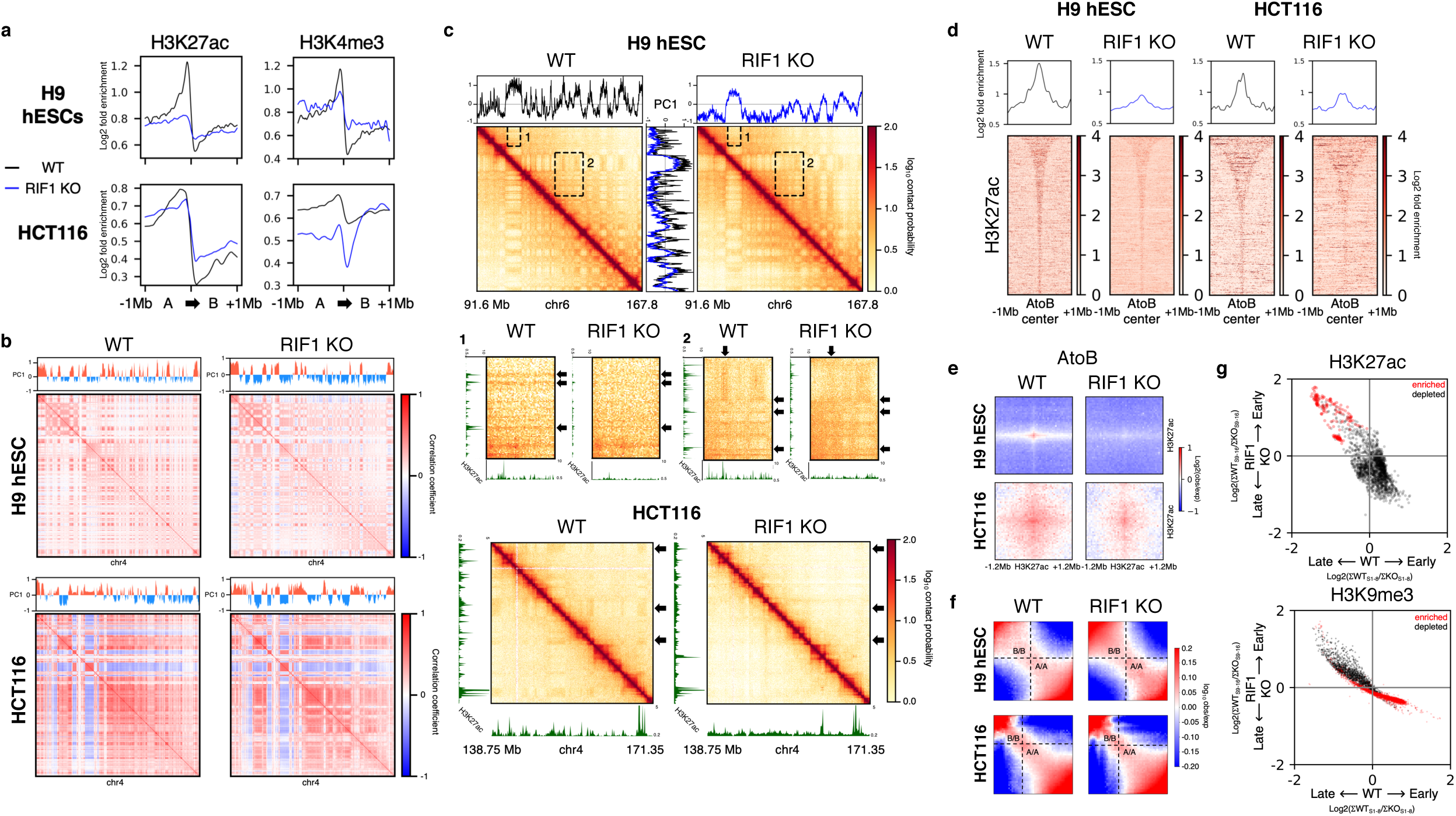
RIF1 KO causes global alterations of compartments and epigenetic state. a, Pile up line plots of indicated histone marks at A to B compartment boundary ±1 Mb in WT (black) and RIF1 KO (blue) HCT116 and H9 hESCs. b, Correlation matrices and PC1 Eigenvector of Chr4 in WT and RIF1 KO H9 hESCs (top) and HCT116 (bottom). c, Example ICE normalized Hi-C contact map of Chr6 91.6-167.8 Mb in H9 hESC (top) and Chr4 138.75-171.35 Mb in HCT116 (bottom) WT and RIF1 KO cells with accompanying PC1 Eigenvector plots. Eigenvector plots are overlaid between contact maps. Below are expanded views of insets 1 and 2 with accompanying H3K27ac ChIP-seq plots. Arrows indicate compartments and ChIP-seq peaks that are lost upon RIF1 KO. d, Fold enrichment signal pile ups for H3K27ac centered on AtoB compartment switching regions ±1 Mb in WT and RIF1 KO cells sorted by size. Line plots represent cumulative signal in WT (black) and RIF1 KO (blue) cells. e, Aggregate Hi-C log2(obs/exp) interactions between H3K27ac peaks within AtoB compartment switching regions ±1.2 Mb in WT and RIF1 KO H9 hESCs. f, Genome wide saddle plots in WT and RIF1 KO cells. g, Scatterplot of the RT values for upregulated (red) and downregulated (black) H3K27ac peaks in WT (x axis) and RIF1 KO (y axis) HCT116 cells calculated from high resolution Repli-seq data.

Both cell lines exhibited genome wide weakening of A/A interactions accompanied by strengthening of B/B interactions (Fig 3f). This was further confirmed when we called statistically significant differential interactions (strengthened and weakened) using diffHiC (*21*). Strengthened interactions were seen predominantly within the B compartment and weakened interactions were concentrated in the A compartment in both cell lines (Fig S6a, b). This is likely the combined effect of strengthened interactions between upregulated H3K9me3 domains within the B compartment (Fig S5f, g) and weakened interactions between downregulated H3K27ac peaks in the A compartment (Fig 3e). Together these data indicate a substantial reorganization of compartments in accordance with epigenome changes in RIF1 KO cells.

TAD boundaries, called by directionality index (DI) (*22*), also changed locally in conjunction with shifts in RT and epigenetic state. In individual cases where a large late replicating domain broke into an unaffected region and an affected region upon RIF1 KO, a new TAD boundary formed in accordance with the new RT boundary demarcated by enriched H3K9me3 showing that changes in chromatin state correlated with loss of RT control can permit the formation of new TAD boundaries (Fig S7a). New TAD boundaries were formed within affected late regions in HCT116, consistent with their earlier RT indices and increased A-compartment association (Fig S7b bottom, c) as early replicating, A compartment chromatin typically contains numerous TADs while late, B compartment chromatin contains fewer TADs (*22*). Reciprocally, boundaries were also lost in regions of later RT (Fig S7b top). Genome wide, the number (∼4000 in H9 hESC and ∼2600 in HCT116) and positioning of TAD boundaries remained similar between WT and RIF1 KO (Fig S7d), while the strength of TAD boundaries was increased in RIF1 KO measured by insulation score (Fig S7d, e, f). Consistently, the ratio of intra-TAD/inter-TAD interactions was increased in RIF1 KO, particularly in HCT116, (Fig S7g) suggesting overall TAD strengthening. ChIP-seq of the essential TAD protein cohesin’s subunit Rad21 revealed largely unaffected binding with a small number of up and down regulated peaks in both cell lines (Fig S8a). Focusing on the interactions at common, up, and down regulated Rad21 peaks, we found that boundaries were strengthened around common and up regulated peaks in both cell lines (Fig S8b). In HCT116, boundaries were strengthened even around downregulated peaks (Fig S8b left) suggesting changes in TAD boundaries were not due to changes in Rad21 peak intensity. In H9 hESCs, we observed weakening of boundaries around the small number of downregulated Rad21 peaks (Fig S8b right). However, such boundaries were weak boundaries in WT and constituted a small fraction of total boundaries (Fig S8b right). Overall, these data mechanistically separate RT from TAD formation and support previous work showing essential TAD proteins Rad21 and CTCF are dispensable for global RT control (*23*–*25*).

To investigate if the profound changes in genome organization and epigenetic state seen in RIF1 KO cells affected gene expression we performed RNA-seq on WT and RIF1 KO HCT116 and H9 hESCs. With the threshold for called change set at FDR < 0.1, RIF1 KO caused 2284 genes (1378 upregulated, 906 downregulated) in H9 hESCs and 1737 genes to change expression (818 upregulated, 919 downregulated) in HCT116 (Fig S9a). Gene ontology (GO) analysis revealed expression changes for genes important for cancer progression in HCT116 and developmentally regulated genes in H9 hESCs (Fig S9b). However, H9 hESCs devoid of RIF1 did not have significantly reduced expression of Oct4, Nanog, Sox2, Rex1, or any other key pluripotency factors (Fig S9c), consistent with their ability to self-renew (Fig S1c) with no morphological signs of differentiation. RT and chromatin compartmentalization are correlated with gene expression (*1*), however we observed no universal pattern linking changed gene expression and changed RT or compartmentalization. Neither changes in RT nor compartment switches were able to predict gene expression changes in RIF1 KO cells of either cell type (Fig S9d, e). Loss of late timing was not correlated with increased gene expression nor was loss of earlier timing correlated with decreased expression (Fig S9d). Similarly, shifts from the B to A compartment were not correlated with increased gene expression nor were shifts from A to B correlated with downregulated expression (Fig S9e). However, the WT chromatin context of differentially expressed genes did correlate with expression changes. In H9 hESCs where the epigenetic landscape is plastic (*26, 27*) earlier replicating genes with strong A compartment association in WT cells were likely to be downregulated while later replicating genes with weak A compartment association in WT cells were likely to be upregulated upon RIF1 KO (Fig S9d, f).

In HCT116 cells, only genes in early replicating, A compartment chromatin were differentially expressed (both up and downregulated) and mostly maintained their early replication (albeit more temporally heterogeneous) and A compartmentalization upon RIF1 KO (Fig S9d, e, f, g). Conversely, specific chromatin modifications were more correlated to gene expression changes upon RIF1 loss than changes in RT or 3D genomic organization. Differentially expressed genes showed cell type specific changes in the distribution of specific histone modifications around their TSSs that correlated with expression changes (Fig S9h). In H9 hESCs H3K27me3 was high in WT cells around TSSs of upregulated genes and decreased upon RIF1 KO while levels of H3K27me3 remained constantly low around downregulated genes (Fig S9h, left). WT H3K9me3 levels were low around both up and downregulated genes and didn’t change significantly upon RIF1 KO (Fig S9h, left). Both up and downregulated TSSs showed similar changes to the active marks that reflected the global trend of change. H3K27ac was diminished and H3K4me3 remained largely constant (Fig S9h, left). Taken together, random forest regression feature importance analysis showed H3K27me3 to be most important in predicting gene expression changes in H9 hESCs (Fig S9i). As most genes that were differentially expressed in HCT116 were associated with the A compartment and early replication in WT cells, these TSSs showed changes in histone mark distribution similar to changes in the A compartment as a whole (Fig 2j bottom, Fig 3a bottom). H3K27ac was diminished while H3K27me3 levels remained constant at both up and down regulated genes (Fig S9h, right). However, changes in H3K4me3 levels did correlate with expression changes (Fig S9h, right). In summary, we found changes in histone modifications around the TSS were considerably more predictive of gene expression changes than changes to RT and genome compartmentalization.

Temporal separation of the genome into early and late replicating chromatin is thought to help coordinate chromatin states after the passage of the replication fork where early replicating chromatin is likely to be assembled into open euchromatin while late replicating chromatin is likely to be assembled into heterochromatin (*5*). However, there has been no evidence for this on native mammalian chromosomes due to the prior inability to manipulate RT. We found that loci depleted for H3K27ac were in genomic bins that became later replicating on the population level as a result of RIF1 KO (Fig 3g top). Reciprocally, those few peaks that became more enriched for H3K27ac in RIF1 KO were in bins that became earlier replicating (Fig 3g top). A similar correlative change was seen at H3K9me3 peaks where loci that became depleted for H3K9me3 became earlier replicating while enriched peaks became later replicating (Fig 3g bottom). This correlation is consistent with the model that RT plays a role in establishing the characteristics of nascent chromatin (*5*). Our observation that global histone modification levels are not affected by RIF1 KO (Fig S5d, i) also supports this model as altered RT would predict redistribution of chromatin modifications rather than global gain or loss of histone marks. Furthermore, RIF1 KD in H9 hESCs showed mild disruptions to RT (Fig S2b, c), but no changes in genome compartmentalization (Fig S2d, e, f) indicating that RIF1’s primary role is to control RT and disruptions to epigenetic state and genome architecture are downstream and manifest only when RT disruption are severe.

To more directly address RIF1’s primary mode of action we took advantage of the auxin inducible degron (AID) system (*28*) in HCT116 cells to rapidly degrade GFP tagged RIF1. After 24 hours of RIF1 degradation (Fig 4a) constituting ∼1 cell cycle the RT phenotype observed in RIF1 KO HCT116 cells was fully recapitulated while changes to genome organization were not as striking as RIF1 KO. Genome wide RT defects assayed by E/L Repli-seq (Fig 4b, c) showed RT values centered around zero log2E/L and the retention of a subset of late replicating regions in RIF1-AID cells similar to RIF1 KO (Fig 4b, c). However, RIF1-AID depleted cells showed few compartmentalization switches (Fig 4d) and genome-wide intra- and inter-compartment interactions were similar to RIF1-AID control cells (Fig 4e). Moreover, Euclidean distance measurements between RIF1-AID depleted cells and either RIF1-AID control cells or RIF1 KO cells showed genome wide RT values of RIF1-AID depleted cells were closer in Euclidean distance and thus more similar to RIF1 KO cells. By contrast, genome wide Hi-C log2(observed/expected) values of RIF1-AID depleted cells were more similar to RIF1-AID control cells (Fig 4f). Furthermore, RIF1-AID depleted cells did not show increased interactions between upregulated H3K9me3 domains called in RIF1 KO HCT116 cells (Fig 4g, h; compare to Fig 2j and Fig S5d respectively) nor did these cells lose interaction between depleted H3K27ac peaks called in RIF1 KO HCT116 cells (Fig 4i; compare to Fig 3e) suggesting that the epigenetic chromatin state and thus genome compartmentalization are maintained after 24 hours of RIF1 depletion despite a nearly complete effect on RT. These data provide further evidence that RIF1’s primary role is to control RT and suggest that several cell cycles of severely disrupted genome wide RT is required to significantly affect genome compartmentalization, placing altered RT upstream of the other changes.

**Figure 4.**
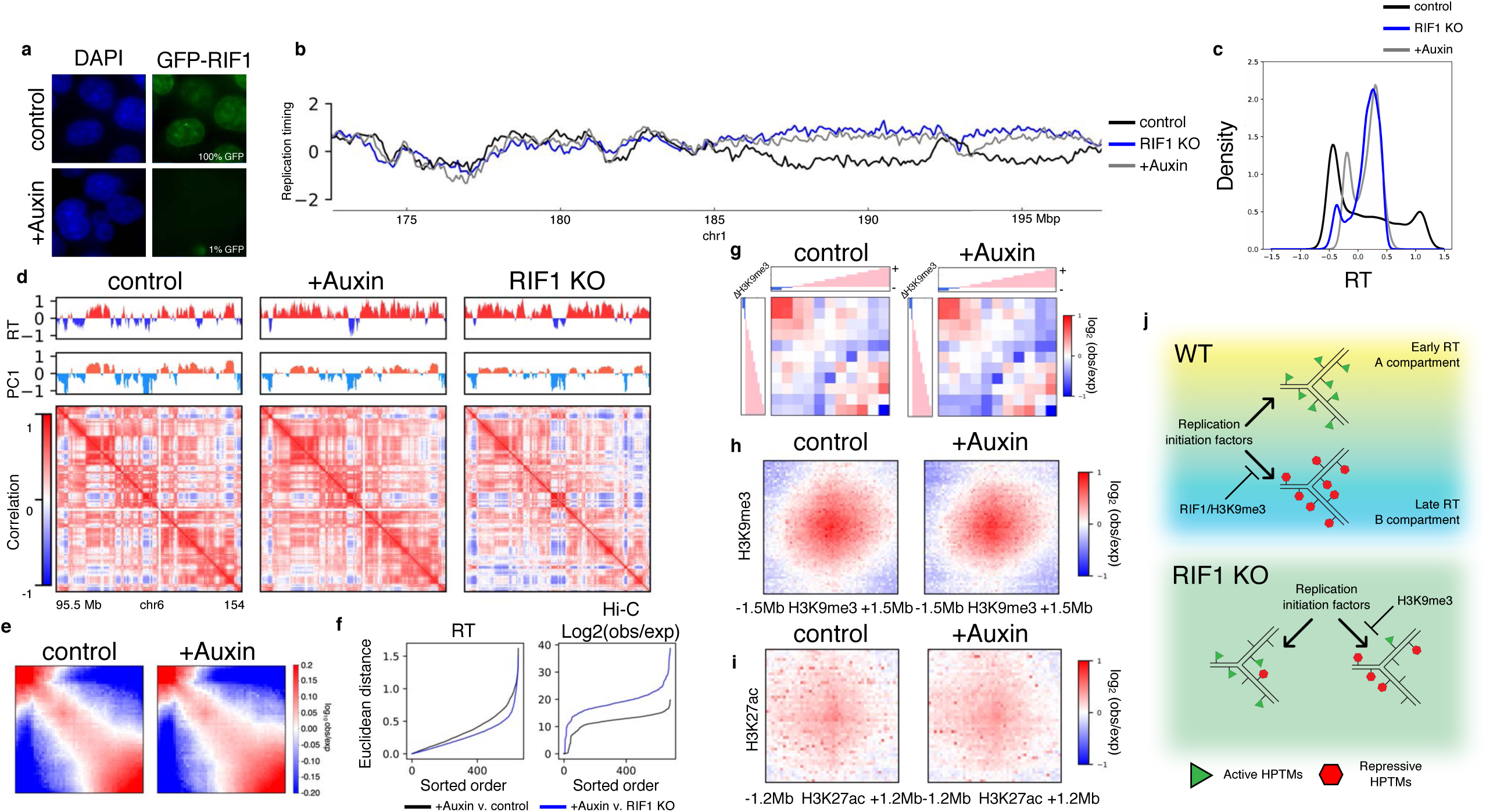
RT effects precede compartment changes. a, Images of HCT116-mAID-mClover nuclei after 24 hrs with DMSO (top) or 500uM Auxin (bottom). b, E/L Repli-seq plots of Chr1 172.6-197.6 Mb in HCT116 RIF1-mAID-mClover cells plus 24hrs DMSO (black), plus 24hrs Auxin (grey) and HCT116 RIF1 KO (blue). c, log2 (E/L) RT probability density plots of HCT116-mAID-mClover cells plus 24hrs DMSO (black), plus 24hrs Auxin (grey) and RIF1 KO (blue) HCT116 cells. d, E/L log2(RT) (top row), PC1 Eigenvector (middle row) and correlation matrices (bottom row) of Chr6 95.5-154 Mb in HCT11-mAID-mClover cells plus 24hrs DMSO (left), plus 24hrs Auxin (middle) and RIF1 KO HCT116 (right). e, Saddle plots showing *cis-*compartmental contacts of HCT11-mAID-mClover cells plus 24hrs DMSO (left) and plus 24hrs Auxin (right) binned into 50 segments of increasing PC1 Eigenvector values. f, Ordered Euclidian distance measurements for genome wide RT (left) and genome wide log2(obs/exp) (right) between HCT116 RIF1-mAID-mClover plus 24hrs Auxin and either HCT116 RIF1-mAID-mClover control cells (black) or HCT116 RIF1 KO cells (blue). g, Log2(obs/exp) aggregate interactions between B compartments in HCT116 RIF1-mAID-mClover cells plus 24hrs DMSO (left) and plus 24hrs Auxin (right). The interactions were binned into 11 equal segments ranked by delta H3K9me3 between WT HCT116 and RIF1KO HCT116. h, Aggregate Hi-C log2(obs/exp) interactions ±1.5 Mb between H3K9me3 domains as defined in RIF1 KO HCT116 cells in HCT116 RIF1-mAID-mClover cells plus 24hrs DMSO (left) and plus 24hrs Auxin (right). i, Aggregate Hi-C log2(obs/exp) interactions ±1.2 Mb between H3K27ac peaks within AtoB compartment switching regions as defined in RIF1KO HCT116 cells in HCT116 RIF1-mAID-mClover cells plus 24hrs DMSO (left) and plus 24hrs Auxin (right).j, A model figure depicting the role of replication timing in maintaining epigenetic status and compartmentalization. In WT cells temporal segregation of replication contributes to the likelihood of nascent chromatin to the assembled into euchromatin (early replicating) or heterochromatin (late replicating). RIF1 and H3K9me3 maintain this temporal segregation in WT cells and allow for preferential 3D contacts between like chromatin types. In RIF1 KO cells this temporal segregation is majorly disrupted, decreasing the likelihood of nascent chromatin assembly into the correct parental form and disrupting 3D contacts between chromatin.

In conclusion, we have shown that late replicating genome is composed of two types of domains, whose delayed replication is enforced by either H3K9me3 or RIF1. The two types of domains form separate hubs of B compartment interactions. Deletion of RIF1 results in genome wide RT disruption resulting from an increase in cell to cell heterogeneity of replication initiation rather than discrete changes in RT. In both hESCs and HCT116 cells, RIF1 KO leads to widespread aberrant histone modification patterns which correlate with distinct genome-wide changes in 3D genome architecture and gene expression. We further show that RT abrogation precedes changes in compartmentalization. We propose that RT changes due to RIF1 KO results in aberrant re-establishment of epigenetic marks at replication forks that causes profound changes in the epigenetic landscape that then alter genome architecture (Fig 4j) rather than RIF1 acting directly to control genome organization as has been previously proposed (*11*). We also show that despite massive epigenetic de-regulation, cells proliferate nearly normally. Since Rif1 KO mice die in utero after gastrulation (*29*), we speculate that RIF1 is needed to coordinate epigenetic changes during cell fate transitions but it, and the proper regulation of epigenetic state, is dispensable for basic cell growth and proliferation. Further exploration of the effects of RIF1 KO on lineage commitment in hESCs us thus poised to unveil insights into the role of epigenetic stability in early human development. Our work establishes RIF1 as a key regulator of epigenome maintenance through its role in RT establishment. Our experiments exploit this role of RIF1 to provide the first mechanistic evidence linking the RT program with maintenance of the global epigenetic state.

## Supplementary Materials

**Fig. S1. CRISPR/Cas9 RIF1 KO does not cause cell cycle arrest.**

a, RIF1 KO in H9 hESCs was achieved by CRISPR/Cas9 cutting at regions up and down stream of exon 3 in the RIF1 coding sequence. Knock in of an eGFP tag was achieved by CRISPR/Cas9 cutting at a region near the start codon and providing a repair vector containing the eGFP coding sequence and flanking homology arms. RIF1 KO HAP1 cells were purchased from Horizon Discovery (HZGHC000663c010) but had a similar deletion of exon3. b, RIF1 GFP tagging in HCT116 was achieved by CRISPR/Cas9 cutting at regions upstream of exon 2 and downstream of exon 3 and providing a repair vector that contained an eGFP tagged version of exon 2 and a loxP flanked exon 3 as well as a blasticidin resistance gene. Cre expression was then used to remove exon 3. In both cell lines removal of exon 3 caused a premature stop codon to be added to the RIF1 mRNA and thus a nonfunctional, truncated protein produced. c, Immunoblot of RIF1 protein in WT and two KO clones of HCT116 and H9 hESCs. Ponceau S total protein stain shown as loading control. d, 2-dimensional (2D) cell cycle analysis using FACS to detect incorporation of 488 anti BrdU (y axis) and DNA content by propidium iodide (x axis) in WT and RIF1 KO cells. e, Average bar plots of three replicates of 2D FACS analysis for each sample with standard deviation error bars; table of p values by t-test between comparators for each cell line. f, E/L repli-seq plot of chr2 in WT (black) and two clones of RIF1 KO (blue and red) H9 hESC (top). E/L repli-seq plot of chr1 in WT (black) and RIF1 KO (blue) mouse ESCs from (*11*)(bottom).

**Fig. S2. RIF1 KD in H9 hESCs causes intermediate RT changes with no change to genome compartmentalization.**

a, Immunoblot of control and RIF1 shRNA in H9 hESCs. b, Repli-seq data of Chr1 172.6-197.6 Mb in control shRNA and RIF1 shRNA H9 hESCs. Same region as RIF1 KO in Fig 1a. c, Genome wide probability density of E/L Repli-seq RT values in control and RIF1 shRNA H9 hESCs. d, Scatterplot of genome wide PC1 eigenvector values for 250kb bins between RIF1 shRNA H9 hESCs (x axis) and control shRNA H9 hESCs (y axis) (top). Scatterplot of genome wide PC1 eigenvector values for 250kb bins between RIF1 KO H9 hESCs (x axis) and WT H9 hESCs (y axis) (bottom) e, Region Chr3 148.25-151.75 Mb of E/L RT and PC1 eigenvector in shRNA H9 hESCs and RIF1 KO H9 hESCs. f, Log2(obs/exp) aggregate Hi-C interactions centered on genome wide H3K27ac peaks ±1.2 Mb in control shRNA, RIF1 shRNA, and RIF1 KO H9 hESCs.

**Fig. S3. Replication foci patterns are disrupted in RIF1 KO cells.**

a, Example BrdU incorporation patterns of S phase human cells. b, Quantification of percentage of S phase BrdU incorporation patterns in WT and RIF1 KO H9 hESCs and HCT116.

**Fig. S4. High resolution and single cell Repli-seq reveal major RT heterogeneity in RIF1 KO cells.**

a, High resolution Repli-seq pile up plots in WT and RIF1 KO H9 hESCs and HCT116 centered on IZs timing categories called in WT cells ±750 Kb. b, Log10 fold enrichment qPCR results of BrdU incorporated mouse DNA spike in target over BrdU negative mouse DNA spike in background after BrdU immunoprecipitation of each S phase fraction in WT and RIF1 KO H9 hESCs and HCT116 cells to approximate pull-down efficiency. WT and RIF1 KO of both cell lines show consistent and similar pull-down efficiencies. c, High resolution Repli-seq pile ups centered on RT changed regions called in E/L Repli-seq data (top) and RT unchanged regions called in E/L Repli-seq (bottom) in WT and RIF1 KO HCT116 ±750 Kb. d, RT indices calculation method used in e. e, RT indices calculated from high resolution Repli-seq for RT changed regions called in E/L Repli-seq data (top) and RT unchanged regions called in E/L Repli-seq (bottom) in WT and RIF1 KO HCT116. H9 hESCs RT indices not shown because of similarity between all indices in RIF1 KO. Examples H9 hESC RT indices in Fig 2b. f, Genome wide probability density plots of RT changes between WT and KO for each cell line, calculated from high resolution Repli-seq. Dotted lines indicate those regions called as having negative RT_diff_ (green), close-to-zero RT_diff_ (grey), or positive RT_diff_ (red). Table shows percentages of genome within each category. g, Violin plots of genome wide T_width_ of high resolution Repli-seq data in WT and RIF1 KO H9 hESCs and HCT116. h, Boxplot of population-based E/L Repli-seq RT for replicated (red) and unreplicated (grey) bins for each single-cell in both WT (top) and RIF1 KO (bottom) HAP1 ranked between 30 and 70% S-phase progression.

**Fig. S5. RIF1 and H3K9me3 control late replication via independent mechanisms.**

a, Histogram of RIF1 Cut&Run read density versus genome wide RT in H9 hESCs and HCT116. b, Boxplots of size distributions of H3K9me3 peaks or domains in affected and unaffected regions in WT H9 hESCs and HCT116; ***p < 0.0005. c, Scatterplot of H3K9me3 peaks in WT (x axis) and RIF1 KO (y axis) cells. Significant peak differences are those with at least 2-fold difference and an FDR < 0.05. Upregulated peaks colored in green and downregulated peaks colored in cyan. d, Immunoblot of H3K9me3 in WT and RIF1 KO H9 hESCs at three dilutions. Ponceau S total protein stain shown as loading control. e, Venn Diagrams of shared H3K9me3 domains (left) and unaffected RT regions (right) between RIF1 KO HCT116 and H9 hESCs. f, Log2(obs/exp) aggregate Hi-C interactions between H3K9me3 peaks genome wide ±1.5 Mb in WT and RIF1 KO HCT116 and H9 hESCs. g, Log2(obs/exp) interactions between H3K9me3 peaks as called in f over genomic distance in WT (black) and RIF1 KO (red) cells. Log2(obs/exp) interactions between upregulated H3K9me3 peaks (left). Log2(obs/exp) interactions between downregulated H3K9me3 peaks (middle). Log2(obs/exp) interactions between upregulated and downregulated H3K9me3 peaks (right). h, Scatterplot of H3K27ac peak RPM in WT (x axis) and RIF1 KO (y axis) cells. Significant peak differences are those with at least 2-fold difference and an FDR < 0.05. Upregulated peaks colored in green (n = 5697 in RIF1 KO H9 hESCs, n = 3646 in RIF1 KO HCT116) and downregulated peaks colored in cyan (n = 61617 in RIF1 KO H9 hESCs, n = 21799 in RIF1 KO HCT116). i, Immunoblot of H3K27ac in WT and RIF1 KO H9 hESCs at three dilutions. Ponceau S total protein stain shown as loading control.

**Fig. S6. Intra B compartment interactions are strengthened and intra A compartment interactions are weakened in RIF1 KO cells.**

a, 2-D histogram of PC1 eigenvector values of diffHi-C points of downregulated (left) and upregulated (right) Hi-C interactions in H9 hESCs. b, 2-D histogram of PC1 eigenvector values of diffHi-C points of downregulated (left) and upregulated (right) Hi-C interactions in HCT116 cells.

**Fig. S7. TAD positions are maintained, and boundaries are strengthened in RIF1 KO cells.**

a, ICE normalized region at Chr2 75.2-85.45 Mb showing a TAD boundary formation at a region of RT and H3K9me3 change in HCT116. WT RIF1 Cut&Run shown for comparison to RT and H3K9me3 ChIP-seq. Black dotted line indicates TAD boundary in WT cells. Green dotted line indicates new TAD boundary in RIF1 KO cells. b, Log2(obs/exp) interaction pile ups centered at WT and RIF1 KO specific TAD boundaries ±1 Mb in WT and RIF1 KO HCT116 (left). RT indices of WT and RIF1 KO specific TAD boundaries in WT and RIF1 KO HCT116 (right). c, Autocorrelation (y-axis) against lag (x-axis) of insulation scores for unaffected and affected regions in HCT116 WT and RIF1KO. Higher autocorrelation indicates more similarity between insulation scores within the region of interest, therefore fewer boundaries where insulation score increases. Lower autocorrelation indicates more dissimilarity, therefore potentially more boundaries. d, Insulation score pile ups centered on all TAD boundaries genome wide ±1 Mb in WT and RIF1 KO H9 hESCs and HCT116. Line plots represent mean scores in WT (black) and RIF1 KO (red). e, Log2(obs/exp) interaction pile ups centered on TAD boundary ±1 Mb in WT and RIF1 KO H9 hESCs and HCT116. f, Log2(obs/exp) interaction pile ups centered on TAD center ±1 Mb in WT and RIF1 KO H9 hESCs and HCT116. g, Boxplots of log2(intra/interTAD interactions) of A compartment TADs (blue) and B compartment TADs (grey) in WT and RIF1 KO H9 hESCs and HCT116.

**Fig. S8. TAD boundary strengthening is not caused by changes in Rad21 binding in RIF1 KO cells.**

a, Scatterplots of Rad21 ChIP-seq peak read counts in WT (x axis) versus RIF1 KO (y axis). Significant up or down regulated peaks are those with at least 2-fold difference and a FDR < 0.05. Common peaks colored in black, upregulated peaks colored in green, and downregulated peaks colored in cyan. b, Log2(obs/exp) interaction pile ups centered at Rad21 peaks that are common (top), downregulated (middle), or upregulated (bottom) ±600kb between WT and RIF1 KO cells in HCT116 (left) and H9 hESCs (right).

**Fig. S9. Histone modifications, not RT or compartments, correlate with gene expression alterations in RIF1 KO.**

a, Volcano plots of gene expression changes upon RIF1 KO in H9 hESC (left) and HCT116 (right). Gene expression changes with a corrected p value < 0.1 were called as differential expression events (up-regulated genes: red, down-regulated genes: green, genes with non-significant changes: grey). b, GO analysis dot plot of differentially expressed genes in RIF1 KO H9 hESCs and HCT116 where the sizes of dots are proportional to the number of genes and the color coding indicates the direction of expression change (red = activated, blue = repressed). c, Heat maps of regularized log2(count) – regularized log2(mean) of all genes in WT and two RIF1 KO clones in both cell lines. Top row contains significantly upregulated genes. Middle row contains significantly downregulated genes. Bottom row contains genes with no significant expression change. Genes in all rows are ranked by relative expression in WT cells. Positions of key pluripotency factors (blue text) and histone modification writers (black text) are indicated. d, Scatterplots of upregulated (red) and downregulated (green) gene RT values in WT (x axis) and RIF1 KO (y axis) H9 hESCs and HCT116. e, Scatterplot of upregulated (red) and downregulated (green) gene PC1 eigenvector values in WT (x axis) and RIF1 KO (y axis) H9 hESCs and HCT116. f, Box plots of distribution of WT RT values at down (green) and up (red) regulated genes in both cell lines. Each point represents an individual gene. g, Box plots of distribution of WT PC1 eigenvector values at down (green) and up (red) regulated genes in both cell lines. Each point represents an individual gene. h, Fold enrichment pile up line plots of indicated histone modification ±5kb around TSS of either up (right) or down (left) regulated genes in WT (black) and RIF1 KO (blue) H9 hESCs and HCT116. i, Feature importance of random forest regression modelling of histone modification in predicting direction of for gene expression changes in H9 hESCs.

## Materials and Methods

### Cell lines

H9 hESCs were grown in feeder free conditions on Geltrex matrix (Thermo Fisher A14133) coated dishes in StemPro (Thermo Fisher A100701) media according to manufacturer’s specifications. hESCs were passaged with ReLeSR (StemCell Technologies 05872) according to manufacturer’s specifications. RIF1 KO H9 lines were established by transfecting two CRISPR/Cas9 plasmids containing separate sgRNAs that would cut upstream and downstream of RIF1 exon 2. Cells were subcloned and screened by PCR for loss of amplification from exon 2. HCT116 cells were grown in McCoy’s 5A media plus 10% FBS and 1% Pen/Strep. HCT116 lines were established by CRISPR/Cas9 knock in of eGFP-RIF1-FLOX construct and blasticidin selection. Selected cells were dissociated to single cells and diluted to 100 cells per 10cm plate and allowed to grow into colonies for 1-2 weeks. Clones with homozygous integration of the eGFP-RIF1-FLOX construct were screened by PCR. RIF1 was deleted from HCT116 eGFP-RIF1-FLOX cell lines by transient transfection with pAAV-Ef1a-mCherry-IRES-Cre (addgene.com Plasmid #55632) and FACS sorting of mCherry positive cells. Positive cells were dissociated to single cells and diluted to 100 cells per 10cm plate and allowed to grow into colonies for 1-2 weeks. Clones with homozygous deletion of RIF1 exon3 were screened by PCR. RIF1-mAID-mClover HCT116 cells were established as in (*30*). HAP1 WT and RIF1 KO cells were purchased from Horizon Discovery (HZGHC000663c010) and grown in IMDM plus 10% FBS and 1% Pen/Strep according to manufacturer’s instructions.

### RIF1 degradation with AID

RIF1-mAID-mClover HCT116 cells were incubated with 2ug/mL doxycycline and 100uM Auxinole (*31*) for 24 hrs to induce OsTIR1 expression and suppress its activity. Auxinole was washed away and fresh media plus 2ug/mL doxycycline plus 500 uM IAA (Sigma I2886) was added for 24hrs to degrade RIF1. Control cells remained in doxycycline and Auxinole for an additional 24hrs. RIF1 degradation was confirmed by microscopy and western blotting.

### 2 fraction RT profiling and analysis

Genome-wide RT profiles were generated as previously described (*12*) with the following modifications: RT datasets were not normalized by quantile normalization since this processing step will result in the enforcement of WT distribution on KO, thereby preventing the detection of any genome-wide RT changes. The Loess smoothed coverages are scaled using robust scaler from the python package scikit-learn (http://scikit-learn.org) instead. The scaler removes median and scales the datasets to their collective inter-quantile range according to:

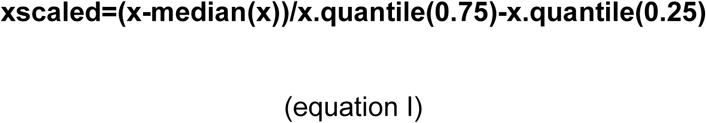

As a result, different datasets with varying dynamic ranges directly comparable to each other. Sex chromosomes were excluded from Repli-seq analysis.

### Replication foci staining

Asynchronous cells were grown on coverslips until 70% confluence and pulsed with BrdU for 30 minutes. Cells were then washed with PBS and fixed in cold 70% EtOH. Chromatin was denatured with 1.5 N HCl 30min RT and washed away. Cells were permeabilized with PBS plus 0.5% Tween20 for 5min RT. Mouse anti-BrdU (BD 55567) (1:50) in PBS plus 10% goat serum was added for 1hr at RT and washed away. Goat 488 anti-mouse (1:100) in PBS plus 10% goat serum was added for 1hr at RT and washed away. Nuclei were stained with DAP1 and cells were imaged on the DeltaVision (GE Life Sciences) microscope. Patterns were categorized as early, middle, late, or blended as in Extended Data Fig 3a.

### High resolution RT profiling and analysis

High resolution Repli-seq was performed and analyzed as previously described (*14*). Briefly, asynchronously growing cells were labeled with BrdU (400 uM) for 30 minutes followed by ethanol fixation. PI staining for DNA content was carried out and 16 fractions of S phase were sorted on the FACS. Gates were set by marking G1 and G2 peaks and dividing S phase into 16 equal fractions between the peaks. At least 80k cells were sorted for each S phase fraction. BrdU incorporated mouse mitochondrial DNA and BrdU negative mouse DNA was spiked into the purified genomic DNA to serve as a BrdU immunoprecipitation control. BrdU incorporated DNA was immunoprecipitated using mouse anti-BrdU (BD 55567). Libraries were prepared for Illumina sequencing with NEBNext Ultra DNA Library Prep Kit for Illumina (E7370). The reads were aligned to hg38 using bowtie2 with parameters --no-mixed --no-discordant --reorder. Replication heatmap matrices were constructed from RPM (read per million) bedgraph files at 50kb window size ranked from S1 to S16. The matrices were Gaussian smoothed with lambda=1 and column-wise scaled. Sex chromosomes were excluded from the heatmap matrix construction.

### Calculating RT indices from high resolution Repli-seq

For each 50kb genomic bin the RT indices computed according to the equation below and graphically illustrated in Supplementary Figure 4b:

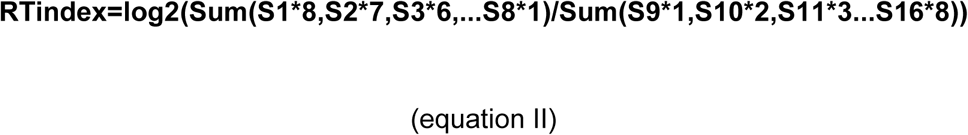

RT_diff_ is computed by subtracting RTindex_KO_ from RTindex_WT_.

### Calling unaffected and affected regions using high resolution Repli-seq

We fitted a Gaussian Mixture model probability distribution on RT_diff_ between WT and KO calculated for all genomic bins. The model is composed of three distribution components: one that contains large negative values representing the bins that on a population level replicate later in WT and earlier in KO, a second distribution of positive values representing bins that on a population level replicate earlier in WT and later in KO, a third one that contains values close to zero representing the bins that on a population level replicate at times statistically indistinguishable from WT. The validity of the model is checked through the minimization of Bayesian information criterion. Continuous 50kb genomic bins that were associated with negative RT indices in WT and assigned to the first distribution were called as affected late regions. Continuous 50kb genomic bins that were associated with negative RT indices in WT but assigned to the third distribution were called as unaffected late regions.

### T_width_ from high resolution Repli-seq

The degree of heterogeneity of RT was estimated by performing a sigmoidal fitting on the column wise cumulative replication percentage as previously described (*14*). Briefly, the sigmoidal function below:

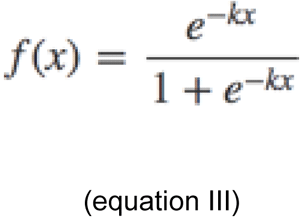

was fitted to bin-wise cumulative replication percentage using curve_fit function in scipy, which aims to minimize the mean squared error. Tolerance is set at 0.0001. T_rep_ used in the timing-variation measurement is f(x) when x = 0.5 which means the genomic bin is 50% replicated in the cell population and T_width_ used in the timing-variation measurement is f(0.75) – f(0.25) which is the time difference between 75% replicated and 25% replicated for any genomic bin.

### Single cell Repli-seq

Single cell RT was performed as previously described (*15*) with slight modifications. Briefly, single cells were sorted from five evenly spaced windows throughout S phase directly into individual wells of a 96 well plate containing single cell lysis buffer and lysed at 50°C for 1 hour followed by 4 minutes at 99°C. Whole genome amplification (WGA) was performed as previously described (*32*) and unique barcodes were added to each individual WGA product followed by purification and pooling for NEBNext Ultra DNA Library preparation.

### Single cell Repli-seq analysis

Single cell Repli-seq fastqs were first demultiplexed according to the barcodes added during library making using custom script. The reads were trimmed 100bp from the 5’ end using CutAdapt (https://github.com/marcelm/cutadapt) and subsequentl aligned to hg38 using bowtie2 with the parameters --no-mixed --no-discordant --reorder. Cells with fewer than 250,000 reads aligned were filtered out. Subsequent analysis was carried out as previously described (*15*). Briefly, mappability correction using G1 cells and smoothing were applied to RPM was calculated in 50kb bins for single cells. Binarization was subsequently carried out on smoothed RPM signal files by applying a 2-component mixed Gaussian model where bins assigned to the distribution with lower mean RPM were binarized to 0 (unreplicated) and those that were assigned to the distribution with higher mean RPM were binarized to 1 (replicated). Heatmaps were generated by sorting according to copy number with the cell with the fewest bins replicated at the top and that with the most bins replicated at the bottom. T_width_ calculation was performed as previously described (*15*). Briefly, for individual bins, an index called ‘time from population average replication’ was calculated, which represents the time in hours passed from the time of replication indicated from E/L repli-seq. A negative number indicates that according to E/L repli-seq the genomic bin should not have undergone replication on a population level whereas a positive number indicates that according to E/L repli-seq the bin should have finished replicating. The 2-D distributions of such indices against % replicated were plotted as heatmaps. A sigmoidal curve was fitted to the distribution as described above in ‘Twidth from high resolution Repli-Seq’.

### ChIP-seq experiments and analyses

ChIP-seq experiments in HCT116 cells and H9 hESCs were carried out as previously described (*33*). After removal of medium, cells were crosslinked in 1% formaldehyde in PBS at room temperature for 10 min and quenched with glycine. PBS rinsed cell pellets were flash frozen in liquid nitrogen and stored at −80°C or continue with cell and nuclear lysis steps. Nuclear lysates were precleared with protein A/G beads followed by incubation with proper antibodies. After washing with high salt buffer, LiCl buffer, and TE, chromatin was eluted and reverse crosslinked. Purified DNA was ethanol precipitated followed by Illumina Truseq library preparation. Libraries for Illumina sequencing were constructed using the following standard protocol. Fragment ends were repaired using the NEBNext End Repair Module and adenosine was added at the 3’ ends of fragments using Klenow fragment (3’ to 5’ exo minus, New England Biolabs), universal adaptors were ligated to the A-tailed DNA fragments at room temperature for 1 h with T4 DNA ligase (New England Biolabs) and amplified with Illumina barcoded primers using KAPA SYBR FAST qPCR Master Mix for 5∼12 PCR cycles to obtain enough DNA for sequencing. Generated libraries were paired-end sequenced on Illumina HiSeq2500 v4. ChIP-seq datasets were aligned to hg38 using bowtie-2 with the parameters --no-mixed --no-discordant --reorder. Resulting bam files for each histone mark were sorted and deduplicated using samtools. Deduplicated bam tools for ChIPed and input libraries were passed onto the peak calling algorithm MACS2 for peak calling and generating fold enrichment signal tracks. The heatmaps centered at features were constructed by filling a matrix where the rows represent individual ChIP-seq fold enrichment signals around individual features.

### RIF1 Cut&Run

Cut&Run was performed as previously described (*16*) with antibody against eGFP in eGFP-RIF1 tagged H9 hESCs and HCT116. Briefly, cells were washed and bound to Concanavalin-A-coated magnetic beads, and permeabilized in wash buffer (20 mM HEPES pH 7.5, 150 mM NaCl, 0.5 mM spermidine and one Roche Complete protein inhibitor tablet per 50 mL) containing 0.03% digitonin and 20 mM EDTA (Dig-wash). Antibody was added to a final concentration of 1:100 and incubated overnight at 4°C. Cells were washed with Dig-wash buffer and incubated with Protein-A-MNase (pA-MN) for 1 hour at 4°C. Cells were washed three times with Dig-wash buffer to remove unbound pA-MN before placing cells in an ice-cold block and activating cleavage with the addition of CaCl2 to 2mM final concentration. Cleavage was stopped by the addition of 2x Stop Buffer (340 mM NaCl, 20 mM EDTA, 4 mM EGTA, 0.05% Digitonin, 0.05 mg/mL glycogen, 5 μg/mL RNase A), and fragments were released by 30 min incubation at 37°C. Supernatant was recovered and DNA was purified with phenol:chloroform extraction and precipitation with ethanol, before being used as input for library preparation.

### Calling H3K9me3 domains

In addition to applying MACS2 (*34*) and epic2 (*35*) on H3K9me3 ChIP-seq datasets to call peaks, we also called large H3K9me3 domain. H3K9me3 fold enrichment was calculated as a log2 ratio over input in 50kb genomic bins. Considering any genomic bin can assume one of two states: inside or outside of a H3K9me3 domain, a two-state hidden markov model was fitted to the fold enrichment distribution with tolerance set at 0.0001. Implementation was carried out in Python using the hmmlearn package (https://github.com/hmmlearn/hmmlearn). Each genomic bin was assigned a state using the Viterbi algorithm. The state associated with the higher signal mean was inside H3K9me3 domain. H3K9me3 domain was therefore defined as continuous genomic bins associated with the state of higher mean signal.

### Hi-C procedure and sequence processing

Two subclones of H9 and HCT116 RIF1 KO cells were prepared and processed for Hi-C as previously described (*36*) using DpnII for digestion. ∼1 billion 50bp paired-end reads were obtained for each subclone for both H9 and HCT116. Reads were processed using HiCUP pipeline available from Babraham Institute (https://www.bioinformatics.babraham.ac.uk). Briefly the reads were truncated at DpnII recognition sites and mapped to hg38 using bowtie2. The uniquely mapped reads were further filtered to remove common Hi-C artefacts e.g. self-ligated fragments and same frag and subsequently deduplicated. Bam files were converted to cooler files using 4DN DCIC utility bam2pairs and cooler load at 5kb, 50kb, 100kb and 250kb. Hi-C maps used in this work were either normalized using iterative correction (ICE) (*37*) or distance normalized as log2 (observed/expected) where expected is computed according to the equation below:

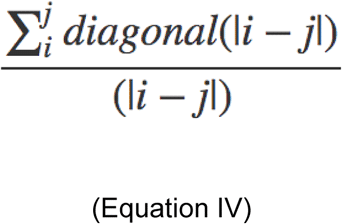

Eigendecomposition is performed using cooltools package and eigenvectors are ranked using gene density.

### TAD calling

TADs and insulating boundaries in this work were called using DI (*22*) and insulation score (*38*) respectively at 50kb bins with a sliding window of 1Mb.

### Calling differential interacting pairs

Differential interacting pairs were called using diffHiC (*21*). Briefly, read counts matrices were generated from bam files resulting from HiCUP for WT and RIF1 KO. Subsequently, edgeR (*39*) was applied to count matrices to estimate technical noise between replicates and call differential pairs by quasi-F test through generalized linear model (glm) fitting. Pairs with >=2-fold difference and <0.05 FDR were called to be up- or down-regulated depending on the sign of fold change.

### RNA-seq

Libraries were prepared with NEBNEXT rRNA Depletion kit (human/mouse/rat) and NEBNEXT Ultra II Directional RNA Library Prep kit for Illumina (New England Biolabs) according to manufacturer’s instructions. Sample input was 900ng total RNA (determined by Qubit RNA HS reagents, Thermo) with a RIN >8 (TapeStation High Sensitivity RNA ScreenTape Assay, Agilent). RNA was fragmented for 15 minutes and libraries were constructed with a 1/5th dilution of NEB adaptor and 12 cycles of PCR amplification with dual-indexing primers. Amplified libraries were initially quantified by Qubit DNA HS reagents and checked for size and artifacts using TapeStation DNA HS reagents. Excess adaptors were removed using an additional 0.9x AMPure bead purification. KAPA qPCR (KAPA Biosystems) was used to determine molar quantities of each library and individual libraries were diluted and pooled at equimolar concentrations. The final library pool was again checked by TapeStation and KAPA qPCR before submission for sequencing. Fastq reads were aligned to hg38 using STAR aligner with the options --quantMode GeneCounts and --bamRemoveDuplicatesType UniqueIdentical turned on. Gene count files were generated using HTseq (https://github.com/simon-anders/htseq) from bam files produced by STAR aligner. DEseq2 (https://bioconductor.org/packages/release/bioc/html/DESeq2.html) was subsequently used to call differentially expressed transcripts using gene count files as input.

### Random Forest Classifier

Random forest classification for differentially expressed genes was performed on arrays where each row represented the TSS of a differentially expressed gene and each column represented the RPM signal of H3K27ac, H3K4me3, H3K27me3 and H3K9me3 according to the algorithm presented in (*40*) using scikit-learn.

## References

1. N. Rhind, D. M. Gilbert, DNA Replication Timing. Cold Spring Harb. Perspect. Biol. 5, a010132 (2013).

2. C. Marchal, J. Sima, D. M. Gilbert, Control of DNA replication timing in the 3D genome. Nat. Rev. Mol. Cell Biol. (2019),, doi: 10.1038/s41580-019-0162-y.

3. N. Reverón-Gómez, C. González-Aguilera, K. R. Stewart-Morgan, N. Petryk, V. Flury, S. Graziano, J. V. Johansen, J. S. Jakobsen, C. Alabert, A. Groth, Accurate Recycling of Parental Histones Reproduces the Histone Modification Landscape during DNA Replication. Mol. Cell (2018), doi: 10.1016/j.molcel.2018.08.010.

4. T. M. Escobar, O. Oksuz, R. Saldaña-Meyer, N. Descostes, R. Bonasio, D. Reinberg, Active and Repressed Chromatin Domains Exhibit Distinct Nucleosome Segregation during DNA Replication. Cell (2019), doi: 10.1016/j.cell.2019.10.009.

5. L. Lande-Diner, J. Zhang, H. Cedar, Shifts in Replication Timing Actively Affect Histone Acetylation during Nucleosome Reassembly. Mol. Cell. 34, 767–74 (2009).

6. M. Hayano, Y. Kanoh, S. Matsumoto, C. Renard-Guillet, K. Shirahige, H. Masai, Rif1 is a global regulator of timing of replication origin firing in fission yeast. Genes Dev. 26, 137–150 (2012).

7. J. M. Peace, A. Ter-Zakarian, O. M. Aparicio, Rif1 regulates initiation timing of late replication origins throughout the S. cerevisiae genome. PLoS One. 9 (2014), doi: 10.1371/journal.pone.0098501.

8. D. Cornacchia, V. Dileep, J.-P. P. Quivy, R. Foti, F. Tili, R. Santarella-Mellwig, C. Antony, G. G. Almouzni, D. M. Gilbert, S. B. C. C. Buonomo, Mouse Rif1 is a key regulator of the replication-timing programme in mammalian cells. EMBO J. 31, 3678–3690 (2012).

9. S. Yamazaki, A. Ishii, Y. Kanoh, M. Oda, Y. Nishito, H. Masai, Rif1 regulates the replication timing domains on the human genome. EMBO J. 31, 3667–3677 (2012).

10. E. Sreesankar, V. Bharathi, R. K. Mishra, K. Mishra, Drosophila Rif1 is an essential gene and controls late developmental events by direct interaction with PP1-87B. Sci. Rep. 5, 10679 (2015).

11. R. Foti, S. Gnan, D. Cornacchia, V. Dileep, A. Bulut-Karslioglu, S. Diehl, A. Buness, F. A. A. Klein, W. Huber, E. Johnstone, R. Loos, P. Bertone, D. M. M. Gilbert, T. Manke, T. Jenuwein, S. C. B. C. B. B. Buonomo, Nuclear Architecture Organized by Rif1 Underpins the Replication-Timing Program. Mol. Cell. 61, 260–273 (2016).

12. C. Marchal, T. Sasaki, D. Vera, K. Wilson, J. Sima, J. C. Rivera-Mulia, C. Trevilla-García, C. Nogues, E. Nafie, D. M. Gilbert, Genome-wide analysis of replication timing by next-generation sequencing with E/L Repli-seq. Nat. Protoc. 13, 819–839 (2018).

13. R. T. O’Keefe, S. C. Henderson, D. L. Spector, Dynamic organization of DNA replication in mammalian cell nuclei: spatially and temporally defined replication of chromosome-specific alpha-satellite DNA sequences. J. Cell Biol. 116, 1095–1110 (1992).

14. P. A. Zhao, T. Sasaki, D. M. Gilbert, High Resolution Mapping of the Temporal Replication Landscape of the Human and Mouse Genome. bioRxiv, 755629 (2019).

15. V. Dileep, D. M. Gilbert, Single-cell replication profiling to measure stochastic variation in mammalian replication timing. Nat. Commun. 9, 427 (2018).

16. P. J. Skene, S. Henikoff, An efficient targeted nuclease strategy for high-resolution mapping of DNA binding sites. Elife (2017), doi: 10.7554/elife.21856.

17. D. Nicetto, K. S. Zaret, Role of H3K9me3 heterochromatin in cell identity establishment and maintenance. Curr. Opin. Genet. Dev. 55, 1–10 (2019).

18. T. Ryba, I. Hiratani, J. Lu, M. Itoh, M. Kulik, J. Zhang, T. C. Schulz, A. J. Robins, S. Dalton, D. M. Gilbert, Evolutionarily conserved replication timing profiles predict long-range chromatin interactions and distinguish closely related cell types. Genome Res. 20, 761–770 (2010).

19. E. Yaffe, S. Farkash-Amar, A. Polten, Z. Yakhini, A. Tanay, I. Simon, Comparative analysis of DNA replication timing reveals conserved large-scale chromosomal architecture. PLoS Genet. 6, 1–12 (2010).

20. E. Lieberman-Aiden, N. L. Van Berkum, L. Williams, M. Imakaev, T. Ragoczy, A. Telling, I. Amit, B. R. Lajoie, P. J. Sabo, M. O. Dorschner, R. Sandstrom, B. Bernstein, M. A. Bender, M. Groudine, A. Gnirke, J. Stamatoyannopoulos, L. A. Mirny, E. S. Lander, J. Dekker, Comprehensive mapping of long-range interactions reveals folding principles of the human genome. Science (80-.). (2009), doi: 10.1126/science.1181369.

21. A. T. L. Lun, G. K. Smyth, diffHic: A Bioconductor package to detect differential genomic interactions in Hi-C data. BMC Bioinformatics (2015), doi: 10.1186/s12859-015-0683-0.

22. J. R. Dixon, S. Selvaraj, F. Yue, A. Kim, Y. Li, Y. Shen, M. Hu, J. S. Liu, B. Ren, Topological domains in mammalian genomes identified by analysis of chromatin interactions. Nature. 485, 376–380 (2012).

23. J. Sima, A. Chakraborty, V. Dileep, M. Michalski, K. N. Klein, N. P. Holcomb, J. L. Turner, M. T. Paulsen, J. C. Rivera-Mulia, C. Trevilla-Garcia, D. A. Bartlett, P. A. Zhao, B. K. Washburn, E. P. Nora, K. Kraft, S. Mundlos, B. G. Bruneau, M. Ljungman, P. Fraser, F. Ay, D. M. Gilbert, Identifying cis Elements for Spatiotemporal Control of Mammalian DNA Replication. Cell. 176, 816–830 (2019).

24. P. Oldach, C. A. Nieduszynski, Cohesin-Mediated Genome Architecture Does Not Define DNA Replication Timing Domains. Genes (Basel). 10, 196 (2019).

25. M. Cremer, K. Brandstetter, A. Maiser, S. S. P. Rao, V. Schmid, N. Mitra, S. Mamberti, K.-N. Klein, D. M. Gilbert, H. Leonhardt, M. C. Cardoso, E. L. Aiden, H. Harz, T. Cremer, Cohesin depleted cells pass through mitosis and reconstitute a functional nuclear architecture. bioRxiv, 816611 (2019).

26. A. Burton, M. E. Torres-Padilla, Epigenetic reprogramming and development: A unique heterochromatin organization in the preimplantation mouse embryo. Brief. Funct. Genomics (2010), doi: 10.1093/bfgp/elq027.

27. R. Strauss, P. Hamerlik, A. Lieber, J. Bartek, Regulation of stem cell plasticity: Mechanisms and relevance to tissue biology and cancer. Mol. Ther. (2012),, doi: 10.1038/mt.2012.2.

28. K. Nishimura, T. Fukagawa, H. Takisawa, T. Kakimoto, M. Kanemaki, An auxin-based degron system for the rapid depletion of proteins in nonplant cells. Nat. Methods. 6, 917–922 (2009).

29. S. B. C. Buonomo, Y. Wu, D. Ferguson, T. de Lange, Mammalian Rif1 contributes to replication stress survival and homology-directed repair. J. Cell Biol. 187, 385–98 (2009).

30. L. P. Watts, T. Natsume, Y. Saito, J. Garzón, M. T. Kanemaki, S. Hiraga, A. D. Donaldson, The RIF1-Long splice variant promotes G1 phase 53BP1 nuclear bodies to protect against replication stress. bioRxiv, 859199 (2019).

31. A. Yesbolatova, T. Natsume, K. ichiro Hayashi, M. T. Kanemaki, Generation of conditional auxin-inducible degron (AID) cells and tight control of degron-fused proteins using the degradation inhibitor auxinole. Methods. 164–165, 73–80 (2019).

32. T. Baslan, J. Kendall, L. Rodgers, H. Cox, M. Riggs, A. Stepansky, J. Troge, K. Ravi, D. Esposito, B. Lakshmi, M. Wigler, N. Navin, J. Hicks, Genome-wide copy number analysis of single cells. Nat. Protoc. 7, 1024–1041 (2012).

33. X. Lyu, M. J. Rowley, V. G. Corces, Architectural Proteins and Pluripotency Factors Cooperate to Orchestrate the Transcriptional Response of hESCs to Temperature Stress. Mol. Cell. 71, 940–955.e7 (2018).

34. Y. Zhang, T. Liu, C. A. Meyer, J. Eeckhoute, D. S. Johnson, B. E. Bernstein, C. Nussbaum, R. M. Myers, M. Brown, W. Li, X. S. Shirley, Model-based analysis of ChIP-Seq (MACS). Genome Biol. (2008), doi: 10.1186/gb-2008-9-9-r137.

35. E. B. Stovner, P. Sætrom, epic2 efficiently finds diffuse domains in ChIP-seq data. Bioinformatics (2019), doi: 10.1093/bioinformatics/btz232.

36. S. S. P. Rao, M. H. Huntley, N. C. Durand, E. K. Stamenova, I. D. Bochkov, J. T. Robinson, A. L. Sanborn, I. Machol, A. D. Omer, E. S. Lander, E. L. L. Aiden, A 3D map of the human genome at kilobase resolution reveals principles of chromatin looping. Cell. 159, 1665–1680 (2014).

37. M. Imakaev, G. Fudenberg, R. P. McCord, N. Naumova, A. Goloborodko, B. R. Lajoie, J. Dekker, L. A. Mirny, Iterative correction of Hi-C data reveals hallmarks of chromosome organization. Nat. Methods (2012), doi: 10.1038/nmeth.2148.

38. B. R. Lajoie, J. Dekker, N. Kaplan, The Hitchhiker’s guide to Hi-C analysis: Practical guidelines. Methods (2015), doi: 10.1016/j.ymeth.2014.10.031.

39. M. D. Robinson, D. J. McCarthy, G. K. Smyth, edgeR: A Bioconductor package for differential expression analysis of digital gene expression data. Bioinformatics (2009), doi: 10.1093/bioinformatics/btp616.

40. L. Breiman, Random forests. Mach. Learn. 45, 5–32 (2001).

