## Supplemental Figures for "Replication timing maintains the global epigenetic state in human cells"

### Fig. S1.

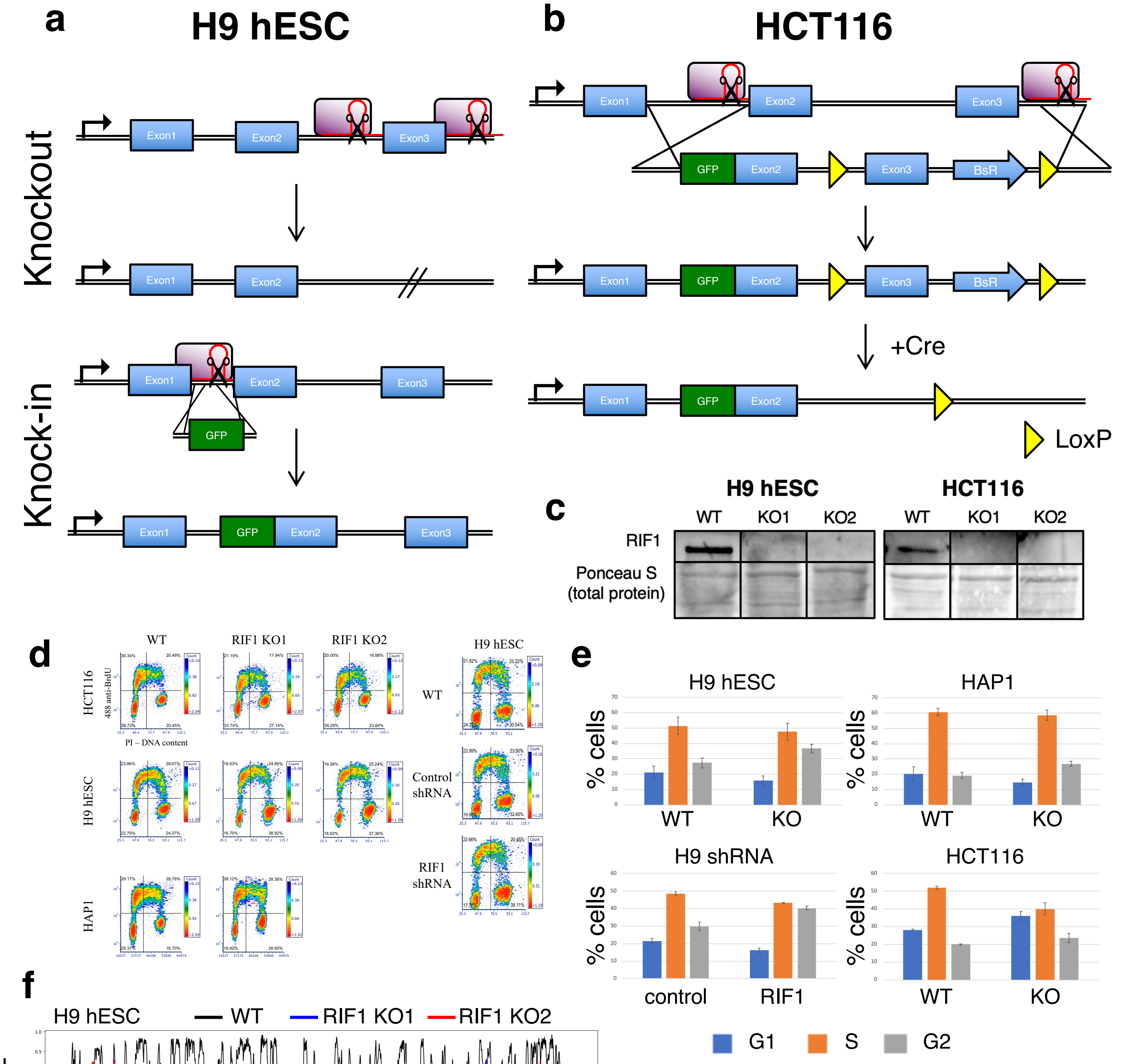


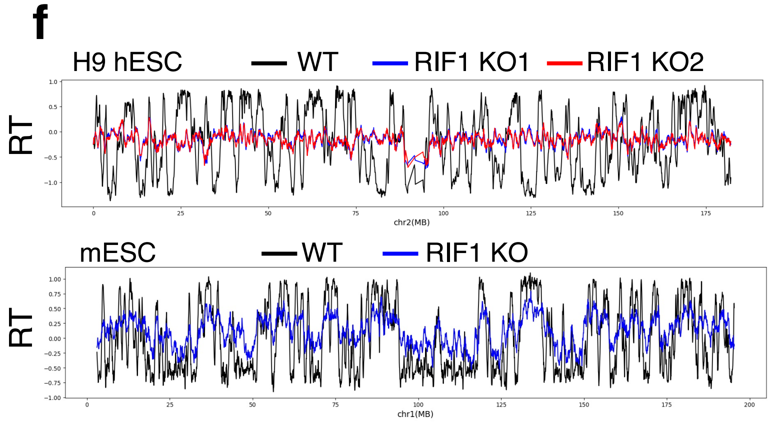


| T test P values | G1 | S | G2 |
|---|---|---|---|
| **H9 hESC KO** | 0.1346 | 0.4508 | 0.0176 |
| **H9 hESC shRNA** | 0.0099 | 0.0009 | 0.0024 |
| **HCT116 KO** | 0.003 | 0.0018 | 0.0808 |
| **HAP1 KO** | 0.1399 | 0.4033 | 0.0081 |

### Fig. S2.

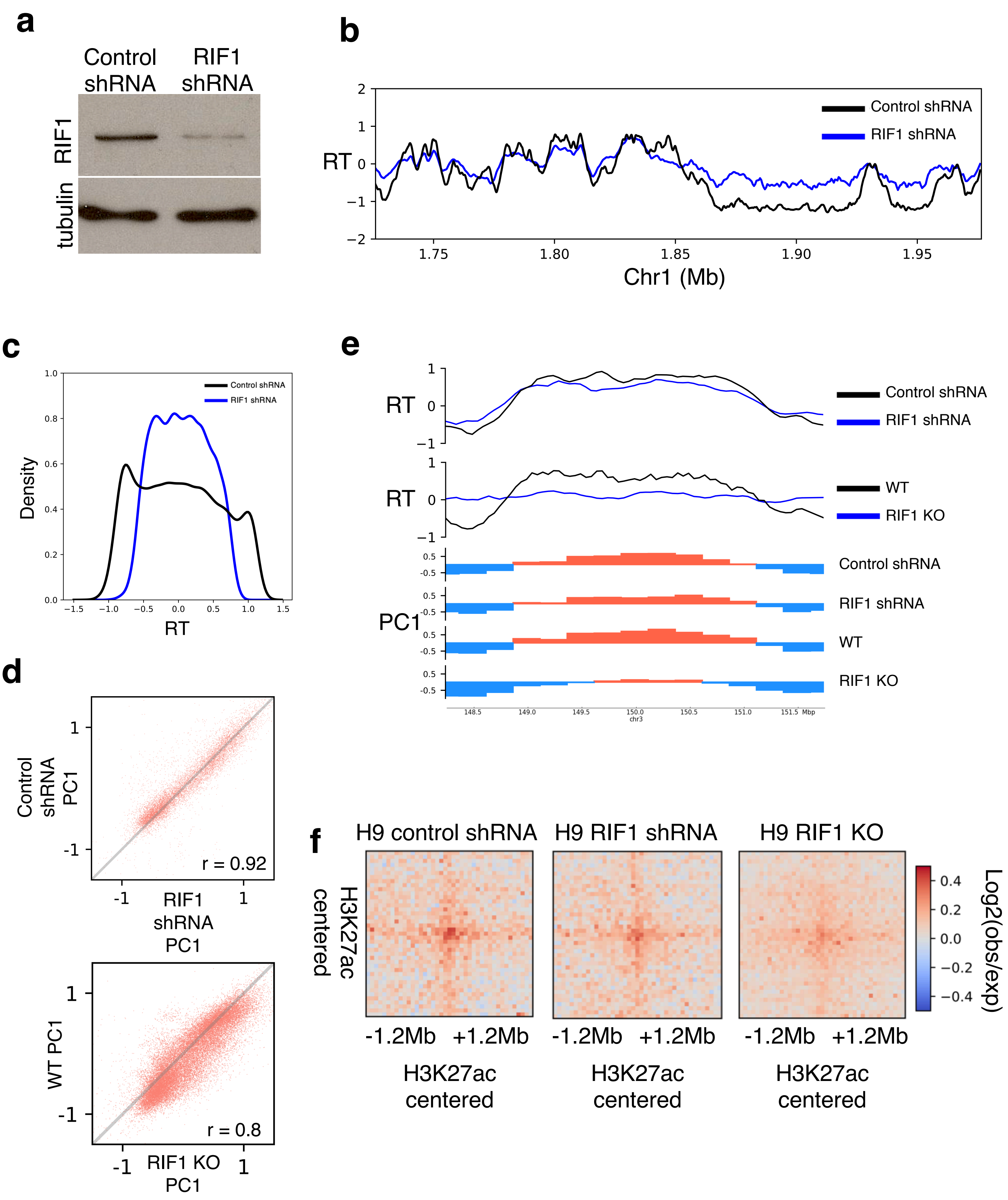
a
Control shRNA
RIF1 shRNA
RIF1
tubulin
b
Control shRNA
RIF1 shRNA
RT
Chr1 (Mb)
c
Control shRNA
RIF1 shRNA
Density
RT
d
Control shRNA PC1
RIF1 shRNA PC1
r = 0.92
WT PC1
RIF1 KO PC1
r = 0.8
e
RT
Control shRNA
RIF1 shRNA
RT
WT
RIF1 KO
PC1
Control shRNA
RIF1 shRNA
WT
RIF1 KO
chr3
Mbp
f
H9 control shRNA
H9 RIF1 shRNA
H9 RIF1 KO
H3K27ac centered
-1.2Mb +1.2Mb
H3K27ac centered
-1.2Mb +1.2Mb
H3K27ac centered
-1.2Mb +1.2Mb
H3K27ac centered
Log2(obs/exp)

**Fig. S3.**

**a**

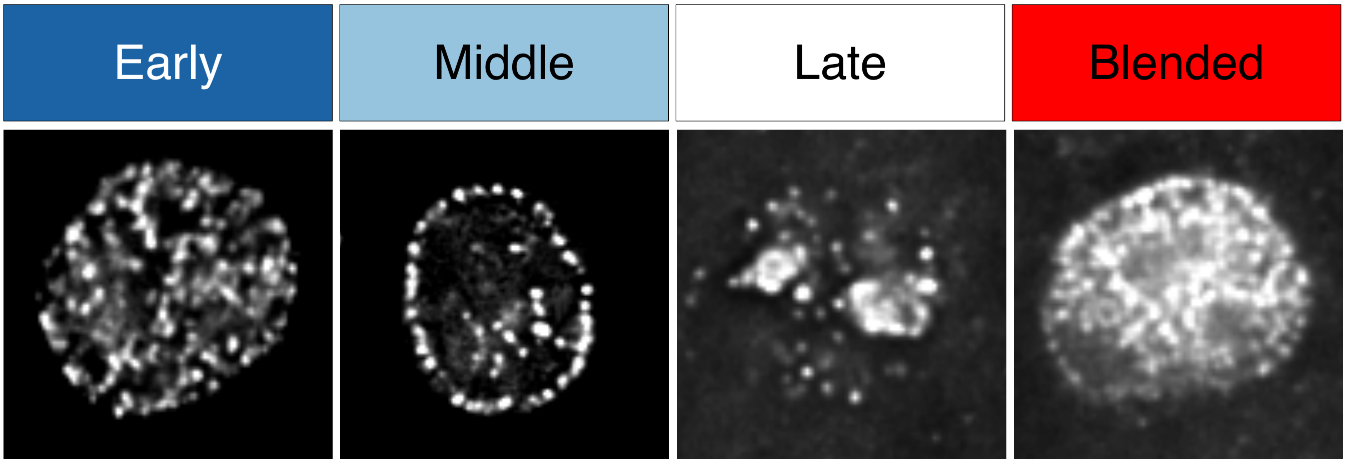
Early
Middle
Late
Blended

**b**

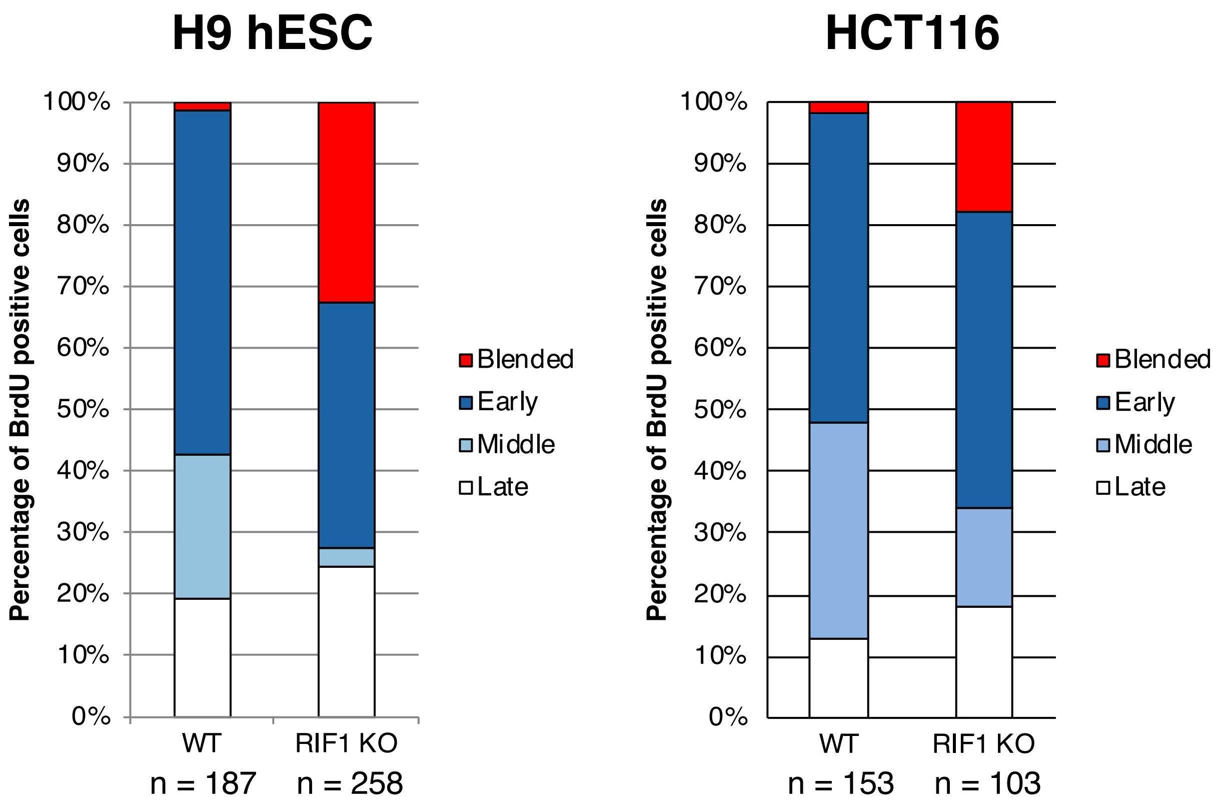
H9 hESC
HCT116
Percentage of BrdU positive cells
100%
90%
80%
70%
60%
50%
40%
30%
20%
10%
0%
WT
n = 187
RIF1 KO
n = 258
WT
n = 153
RIF1 KO
n = 103
Blended
Early
Middle
Late

#### Fig. S4.

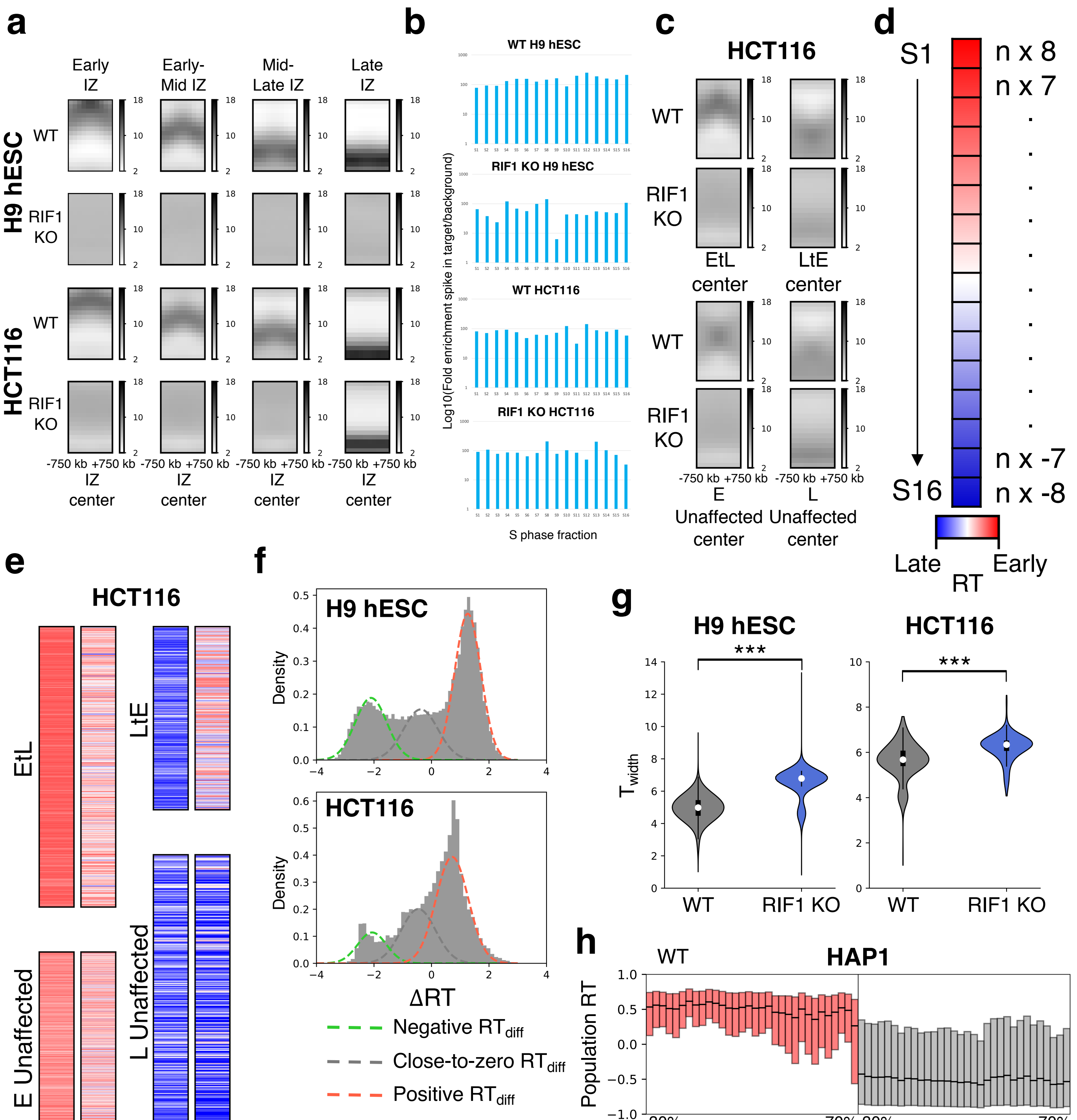


| | H9 hESC | HCT116 |
|---|---|---|
| Negative $RT_{diff}$ | 42% | 30% |
| Close-to-zero $RT_{diff}$ | 14% | 22% |
| Positive $RT_{diff}$ | 44% | 48% |
| Total $RT_{diff}$ | 86% | 78% |

### Fig. S5.

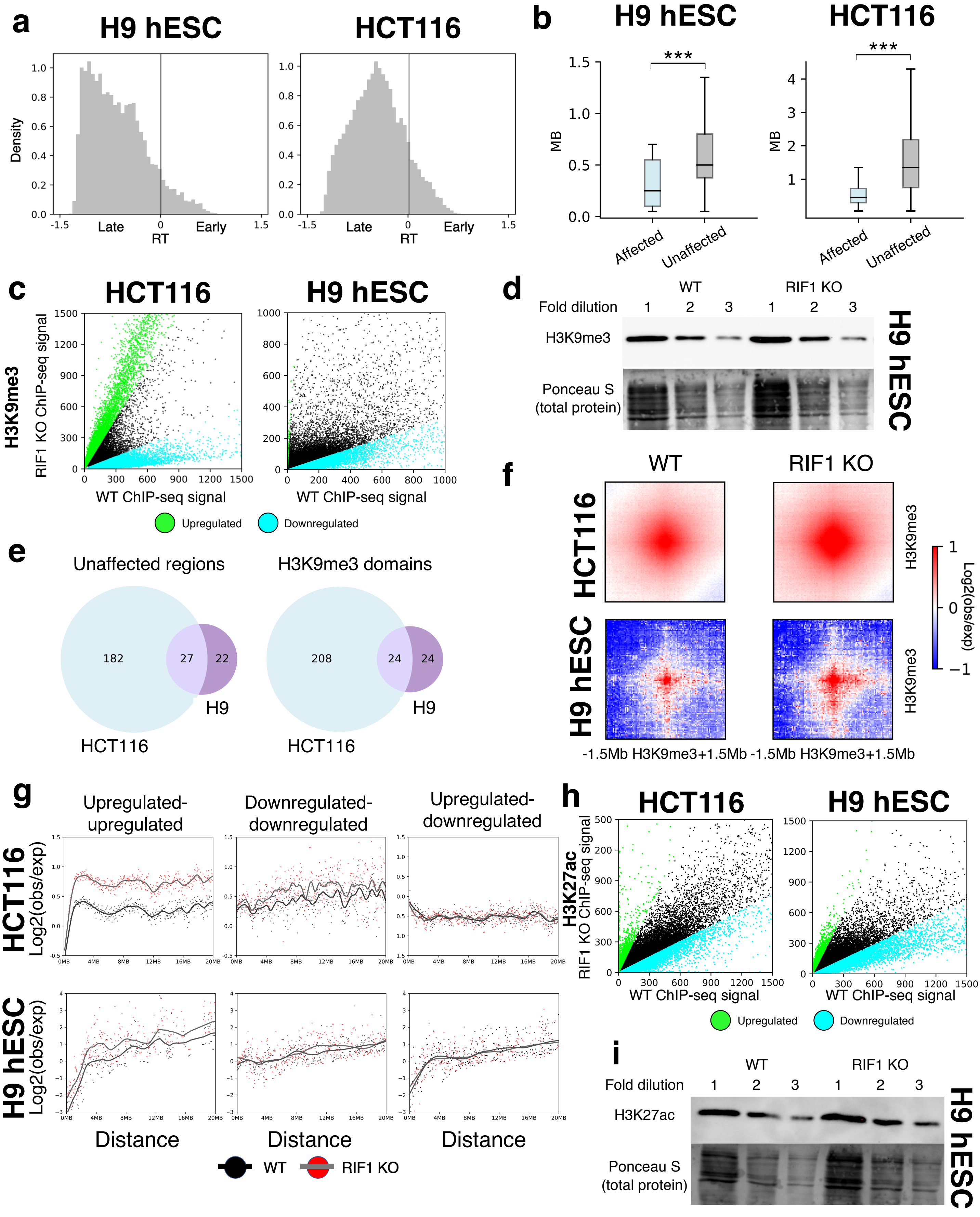
a
H9 hESC
HCT116
Density
Late
Early
RT
b
H9 hESC
HCT116
***
MB
Affected
Unaffected
c
HCT116
H9 hESC
H3K9me3
RIF1 KO ChIP-seq signal
WT ChIP-seq signal
Upregulated
Downregulated
d
WT
RIF1 KO
Fold dilution
1
2
3
H3K9me3
Ponceau S
(total protein)
H9 hESC
e
Unaffected regions
H3K9me3 domains
182
27
22
H9
HCT116
208
24
24
f
WT
RIF1 KO
HCT116
H9 hESC
H3K9me3
Log2(obs/exp)
-1.5Mb H3K9me3+1.5Mb -1.5Mb H3K9me3+1.5Mb
g
Upregulated-
upregulated
Downregulated-
downregulated
Upregulated-
downregulated
HCT116
H9 hESC
Log2(obs/exp)
Distance
WT
RIF1 KO
h
HCT116
H9 hESC
H3K27ac
RIF1 KO ChIP-seq signal
WT ChIP-seq signal
Upregulated
Downregulated
i
WT
RIF1 KO
Fold dilution
H3K27ac
Ponceau S
(total protein)
H9 hESC

**Fig. S6.**

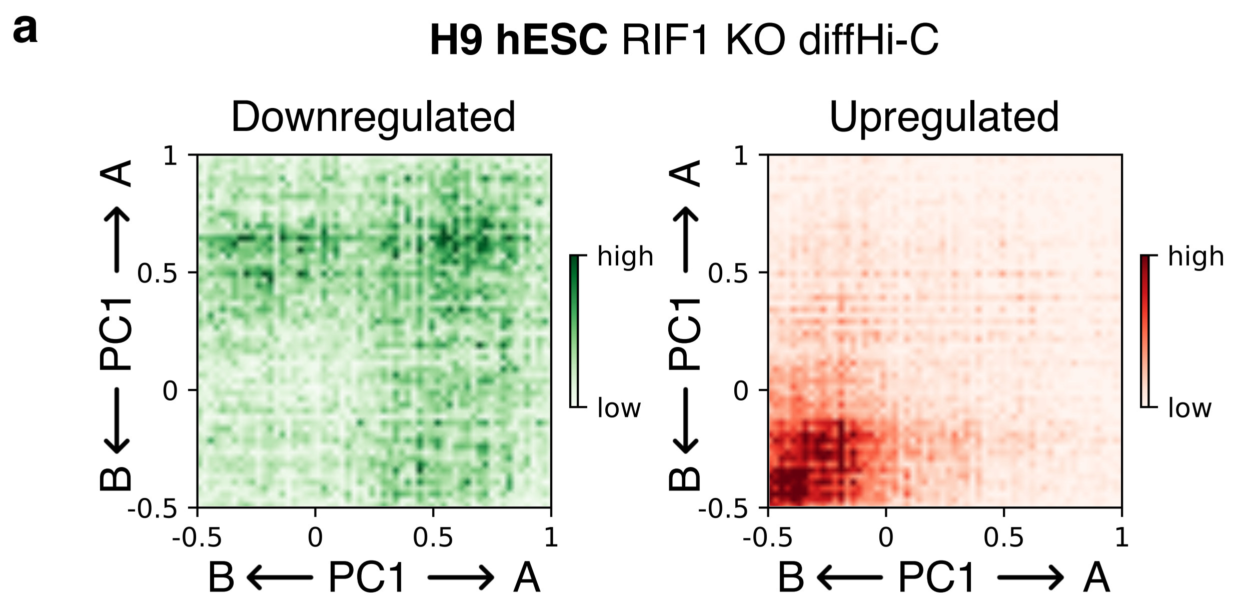
a
**H9 hESC** RIF1 KO diffHi-C
Downregulated
Upregulated
high
low
B ⟵ PC1 ⟶ A
-0.5
0
0.5
1

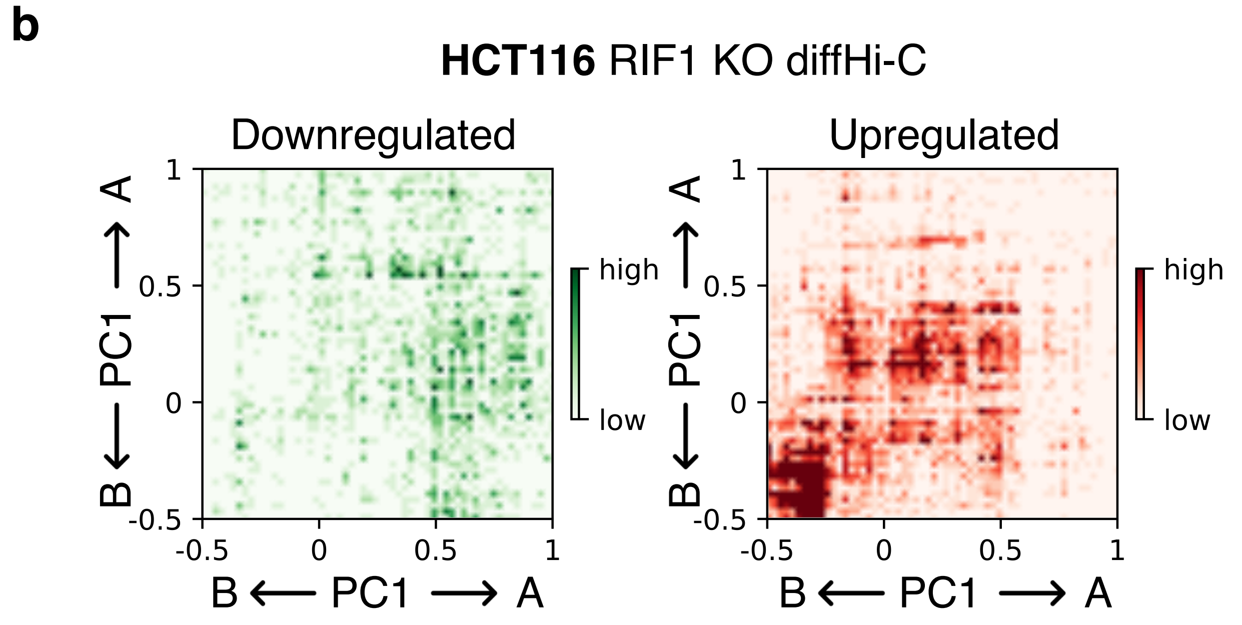
b
**HCT116** RIF1 KO diffHi-C
Downregulated
Upregulated
high
low
B ⟵ PC1 ⟶ A
-0.5
0
0.5
1

### Fig. S7.

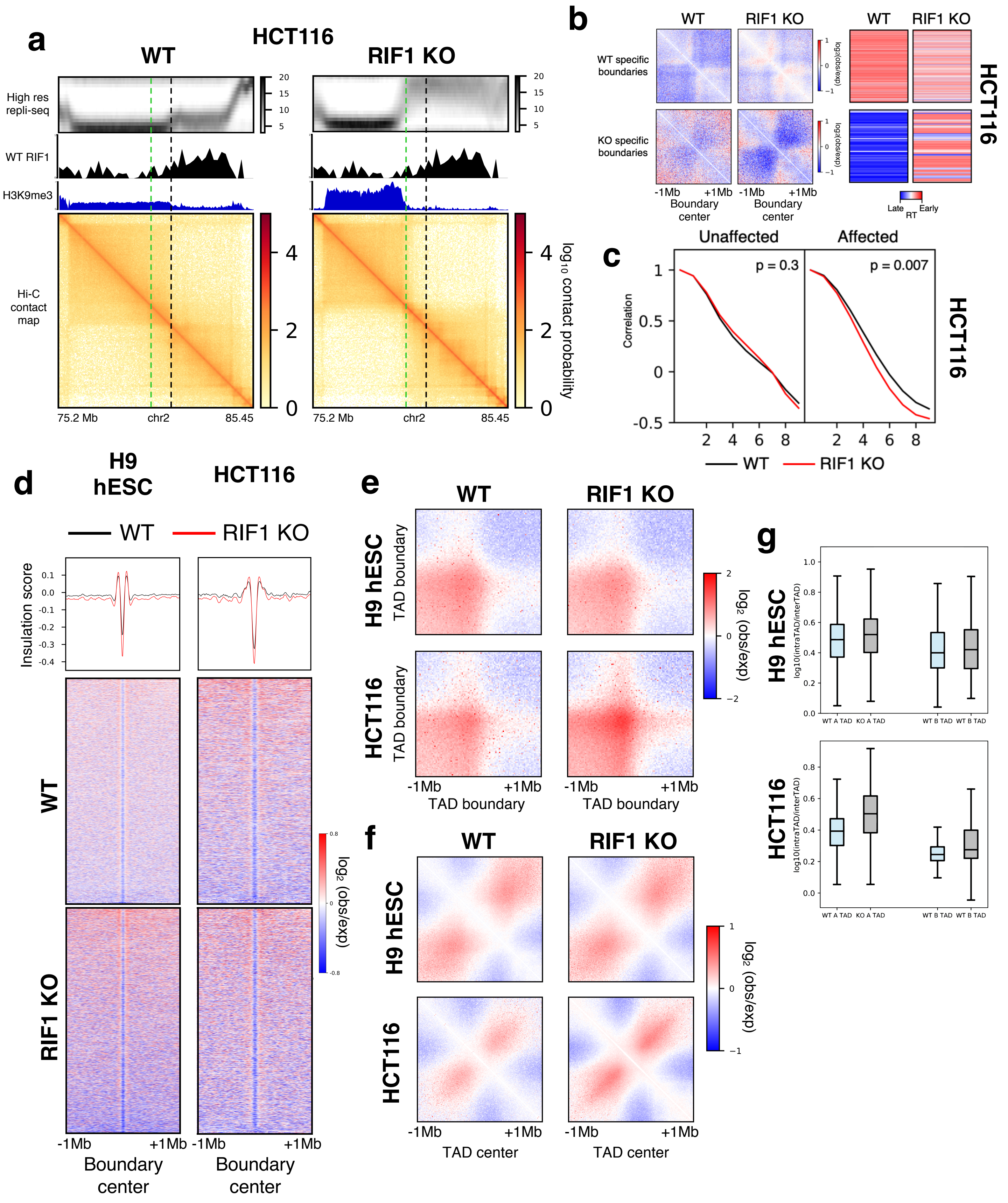
a
HCT116
WT
RIF1 KO
High res repli-seq
WT RIF1
H3K9me3
Hi-C contact map
75.2 Mb
chr2
85.45
log10 contact probability
b
WT
RIF1 KO
WT specific boundaries
KO specific boundaries
-1Mb
+1Mb
Boundary center
log2(obs/exp)
Late
Early
RT
HCT116
c
Unaffected
Affected
p = 0.3
p = 0.007
Correlation
WT
RIF1 KO
HCT116
d
H9 hESC
HCT116
WT
RIF1 KO
Insulation score
Boundary center
log2 (obs/exp)
e
WT
RIF1 KO
H9 hESC TAD boundary
HCT116 TAD boundary
TAD boundary
log2 (obs/exp)
f
WT
RIF1 KO
H9 hESC
HCT116
TAD center
log2 (obs/exp)
g
H9 hESC
HCT116
log10(intraTAD/interTAD)
WT A TAD
KO A TAD
WT B TAD
WT B TAD

### Fig. S8.

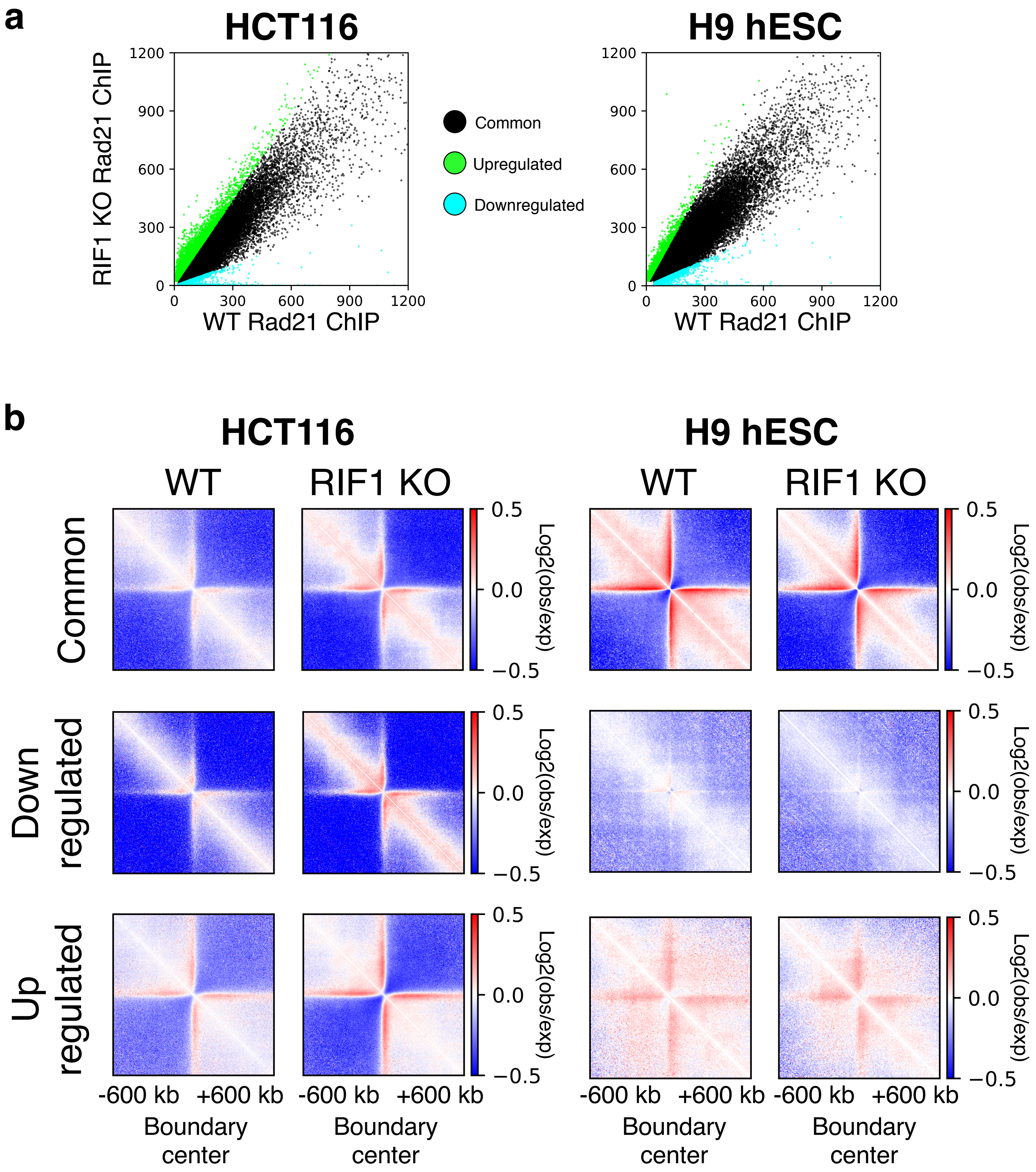
a
HCT116
H9 hESC
RIF1 KO Rad21 ChIP
WT Rad21 ChIP
1200
900
600
300
0
0
300
600
900
1200
Common
Upregulated
Downregulated
b
HCT116
H9 hESC
WT
RIF1 KO
Common
Down regulated
Up regulated
0.5
0.0
−0.5
Log2(obs/exp)
-600 kb
+600 kb
Boundary center

### Fig. S9.

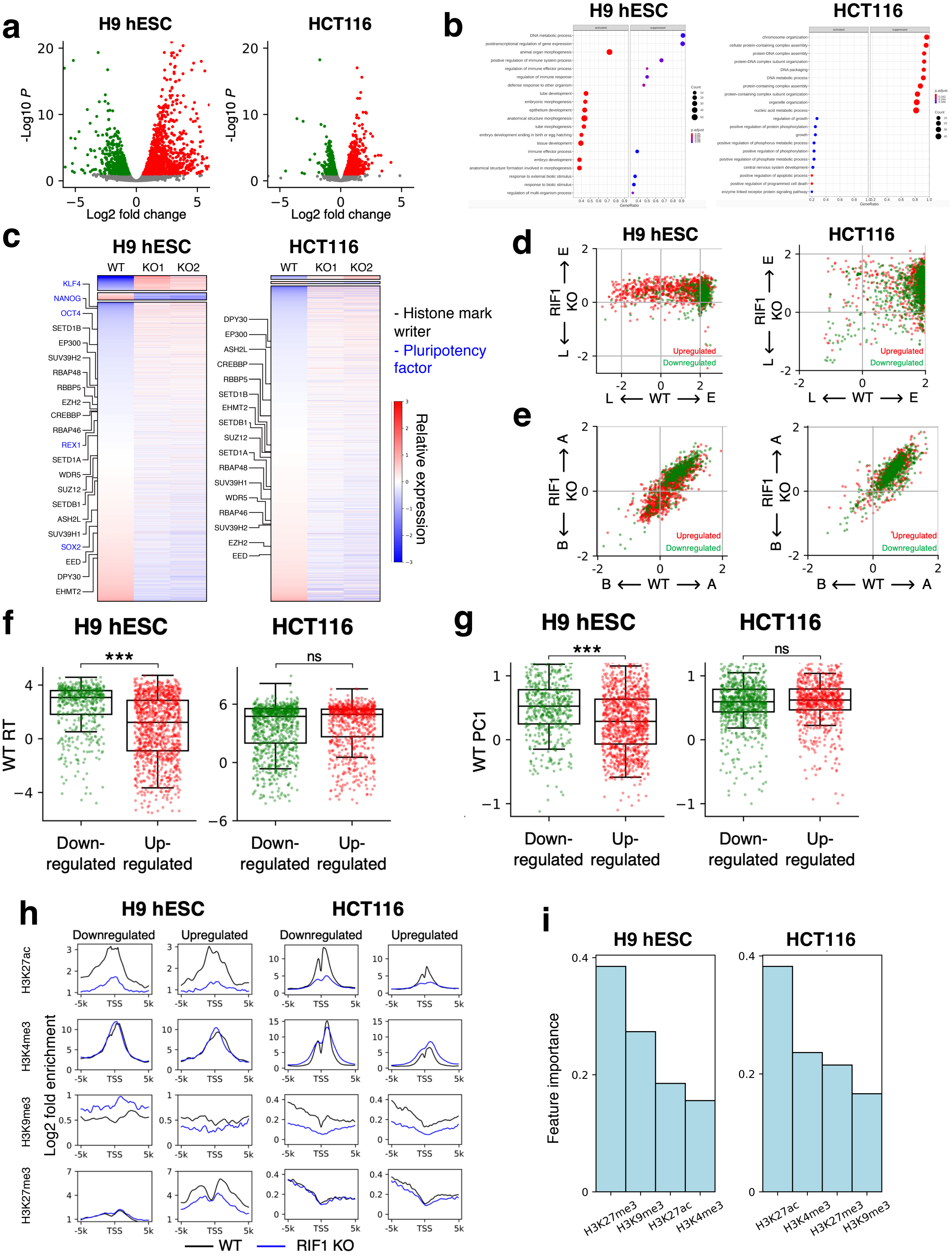
a
H9 hESC
HCT116
-Log10 P
Log2 fold change
b
H9 hESC
HCT116
c
H9 hESC
HCT116
WT
KO1
KO2
- Histone mark writer
- Pluripotency factor
Relative expression
d
H9 hESC
HCT116
RIF1 KO
L ⟵ WT ⟶ E
Upregulated
Downregulated
e
RIF1 KO
B ⟵ WT ⟶ A
Upregulated
Downregulated
f
H9 hESC
HCT116
***
ns
WT RT
Down-regulated
Up-regulated
g
H9 hESC
HCT116
***
ns
WT PC1
Down-regulated
Up-regulated
h
H9 hESC
HCT116
Downregulated
Upregulated
H3K27ac
H3K4me3
H3K9me3
H3K27me3
Log2 fold enrichment
TSS
WT
RIF1 KO
i
H9 hESC
HCT116
Feature importance
H3K27me3
H3K9me3
H3K27ac
H3K4me3
